# Integration of functional immunomonitoring assays with PET/CT scans in TB patients identifies on-treatment biomarkers

**DOI:** 10.64898/2026.05.07.723467

**Authors:** Jingwen Fan, Vincent Rouilly, Munyaradzi Musvosvi, Marie Robert, Chloé Albert-Vega, Vincent Bondet, Annika Jasper, Xiang Yu, Stephanus Malherbe, Raphaël Borie, Nathan Peiffer-Smadja, Karim Sacre, Benjamin Terrier, Gerhard Walzl, Clifton E Barry, Michele Tameris, Thomas J Scriba, Darragh Duffy

## Abstract

Tuberculosis (TB) continues to pose a significant global public health challenge with substantial patient morbidity and mortality. Current TB patient biomarkers lack sufficient resolution to inform treatment response and patient stratification. This necessitates the development of sensitive and reliable host biomarkers. We previously demonstrated the efficacy of TruCulture whole blood stimulation for differentiating asymptomatic TB from active pulmonary TB disease patients in endemic regions. Our systems immunology study expands upon this previous work by evaluating the potential of TruCulture to monitor longitudinal responses to TB treatment in patients from the Predict-TB trial before, during, and after 6 months of antibiotic therapy. We stimulated whole blood from TB patients (n=40) using TruCulture under four conditions (Null, *Mycobacterium tuberculosis*-antigen, LPS, and IL-1β) at baseline (week 0), during treatment (weeks 16 and 24), and one-year follow-up post-treatment (week 72). 20/25 measured cytokines exhibited significant changes throughout treatment, with several continuing to evolve during post-therapy follow-up. Machine learning based analysis identified *Mtb*-Ag-induced IL-1RA (AUC = 0.90, 0.92, 0.95 at weeks 16, 24, 72) and LPS-induced *NLRP3* (AUC = 0.94 at week 16) as the best protein and transcriptional biomarkers for distinguishing treated from untreated patients, strongly implicating the inflammasome response. Combining these results with the extent of lung disease assessed by FDG PET/CT scans, we showed direct disease relevance for these blood-based biomarkers. The identified biomarker profiles hold promise for improving TB patient care through early prediction of treatment responses, real-time therapy monitoring, and informed development of host-directed therapeutic strategies for clinical decision-making.

**Graphical abstract:** Graphical abstract
Predict-TB clinical study overview and summary of TB-specific biomarkers identified from TruCulture whole blood stimulation system.

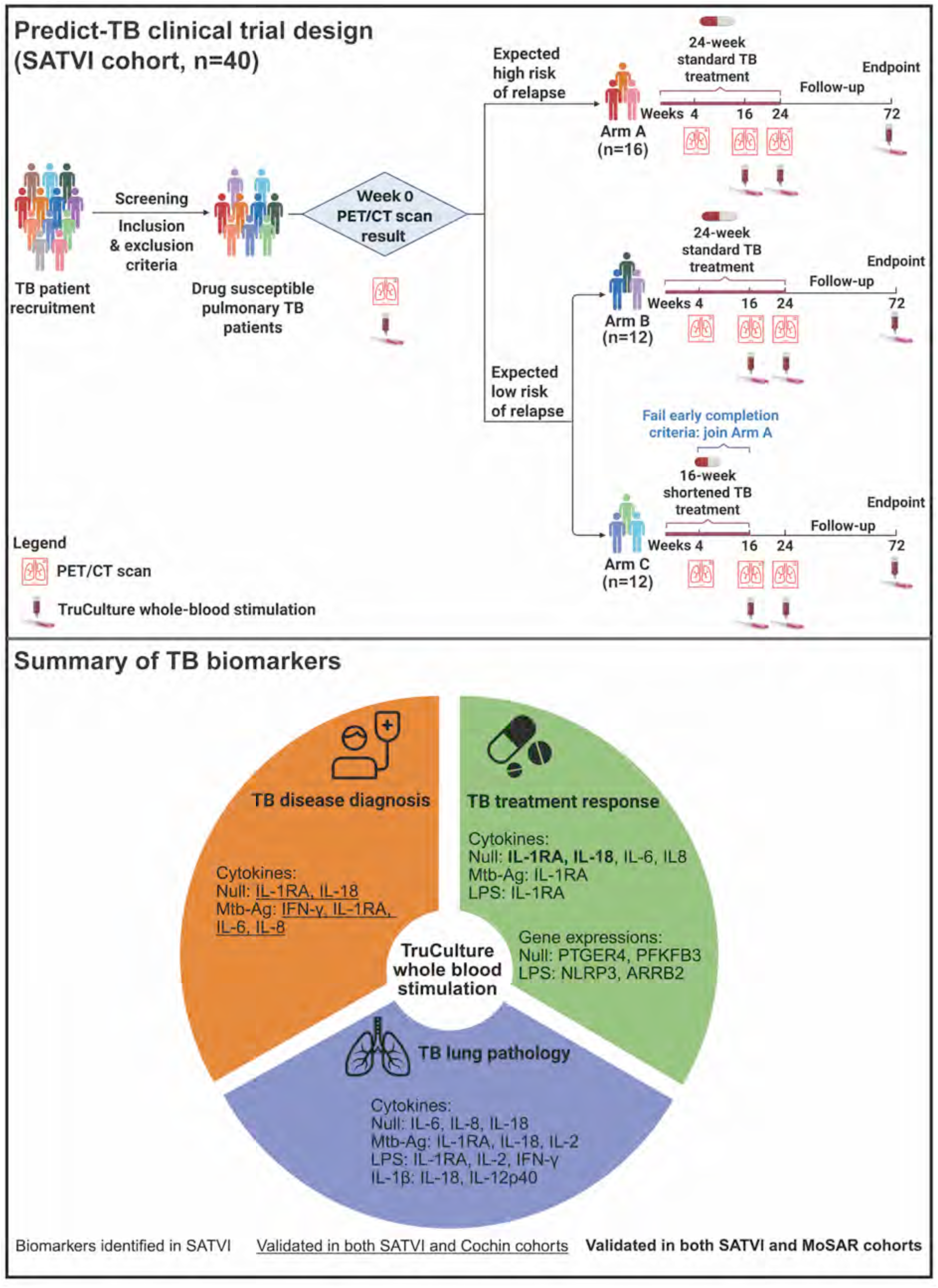

## Introduction

Tuberculosis (TB) remains a major global public health crisis and the leading cause of death from a single infectious agent^1^. An estimated one-quarter of the world’s population has been infected with *Mycobacterium tuberculosis*^2^, with over 10 million new TB cases and more than 1 million deaths occurring each year^1^. The standard first-line treatment regimen for drug- sensitive pulmonary TB requires 6 months of multidrug therapy regardless of TB disease severity. However, emerging evidence suggests that shortened regimens could be sufficient to cure patients with milder TB disease, while some individuals do not achieve cure and may require longer treatment^3,4^. Reducing TB treatment duration without compromising efficacy is essential, particularly in resource-limited, high-TB-burden countries. Most importantly, shortened treatment regimens can improve adherence and mitigate the risk of developing drug-resistant TB, which is a major obstacle to global TB control^1,5^.

A challenge for implementing shortened treatment regimens is the lack of tools to stratify patients into those who require shorter or less intensive therapy and those who may require longer or more intensive therapy^6^. This is complicated by the substantial heterogeneity of TB disease. Over the past decade, there has been growing awareness that TB manifests as a continuous spectrum of disease rather than a binary distinction between latent and active TB^7^. Upon *M. tuberculosis* infection, clinical trajectories vary widely: most infected individuals remain asymptomatic without any evidence of disease or pathology^8^; some clear the infection without treatment^9,10^; some develop asymptomatic disease, while others progress to symptomatic, active TB disease with diverse degrees of severity ^1,6^.

Several bacterial and host-oriented tools have been developed to diagnose and stratify TB patients. Bacterial detection methods are essential for confirming TB disease^1^. Culture-based methods are the gold standard for detecting viable *M. tuberculosis* bacteria and drug resistance on expectorated sputum, but this often requires weeks for results^11^. Automated liquid culture systems such as Mycobacteria Growth Indicator Tube (MGIT) reduce the *M. tuberculosis* detection time to 1-2 weeks in most HIV-negative active TB patients but still lack the rapidity needed for timely clinical decision-making^11^. PCR-based nucleic acid amplification tests (NAAT) such as GeneXpert MTB/RIF enable rapid *M. tuberculosis* detection and drug resistance profiling^11^. However, these bacteriological tools can only be performed in people who can expectorate sputum and provide insufficient resolution for patient stratification. Furthermore, as most patients become bacteriologically negative within weeks of treatment initiation, these tests offer limited utility for ongoing treatment monitoring and cannot inform the risk of TB recurrence. Host-oriented approaches offer complementary information. 2- deoxy-2-[^18^F]fluoro-D-glucose (FDG)–positron emission tomography (PET)/computerized tomography (CT) provides high resolution imaging of thoracic pathology and inflammation and is very informative to assess TB disease severity and guiding treatment decisions^12,13^. Further, CT-detected cavitary lung lesions are a predictor of relapse in shortened TB treatment, although performance is not perfect^13^. However, PET-CT is radioactive, complicated to operate, and very costly, therefore cannot be widely deployed in public health settings to manage TB.

Immunological assays can provide information about the host inflammatory and immune response and can potentially be simple and cost-effective tests. Assays like Tuberculin Skin Test (TST) and QuantiFERON-TB Gold (QFT) measure *M. tuberculosis* immunoreactivity but does not provide information about TB treatment response^10,14^

TruCulture whole blood stimulation is a standardized platform for collection and *ex vivo* stimulation of fresh whole blood^15^. It demonstrates improved reproducibility for immune monitoring compared to peripheral blood mononuclear cell (PBMC)-based assays^16,17^. TruCulture whole blood stimulation has been successfully applied to characterize diverse immune responses in healthy individuals^15,18,19^ and various patient populations^20–23^. This study expands upon our previous work^21^ by evaluating the potential of TruCulture to monitor longitudinal changes in immune responses during and after TB treatment in patients from the Predict-TB trial^24^. In addition, we integrated blood biomarkers with FDG PET/CT results to relate systemic immune responses to lung disease in TB. We further assessed the impacts of demographic and clinical factors such as age, sex, BMI, bacterial burden, previous TB episodes on TB disease progression and treatment responses. Using gene set enrichment analysis (GSEA), we also identified gene expression signatures associated with TB treatment response, TB lung lesions, demographic and clinical factors, providing insights into underlying TB pathogenic mechanisms.

## Results

### TruCulture whole blood stimulation assay qualification

Previous work from our group demonstrated that IFN-γ levels following TruCulture *Mtb*-Ag stimulation best distinguishes active pulmonary TB patients from asymptomatic QFT-positive individuals in a high TB burden setting^21^. To assess the robustness of these previously identified biomarkers for TB diagnosis, we evaluated their repeatability and reproducibility for classifying active TB patients and asymptomatic QFT-positive controls (Fig. S1A). *Mtb*-Ag-induced IFN-γ showed high consistency (91% repeatability; 85% reproducibility). In addition, other biomarkers identified in previous work^21^, including unstimulated (Null) IL-1RA and IL-18, and *Mtb*-Ag stimulation induced IL-6, IL-8, and IL-1RA also demonstrated good performance, with repeatability and reproducibility non-inferior to *Mtb*-Ag-induced IFN-γ (Fig. S1B).

### Study design and patient characteristics

To assess the capability of TruCulture whole blood stimulation to monitor TB treatment responses, we studied 40 HIV-negative participants with drug susceptible, pulmonary TB enrolled in the Predict-TB clinical trial (clinicaltrials.gov NCT02821832)^24^. Following baseline FDG PET/CT evaluation^13^, TB patients were stratified by predicted relapse risk and assigned to three treatment arms. High-risk patients received 24-weeks of standard treatment (Arm A, n=16). Low-risk patients were randomized to either 24-week (Arm B, n=12) or 16-week (Arm C, n=12) regimens^13^. Baseline characteristics were balanced across study arms except for radiological lung lesion severity^13^ (Table 1). Patients in Arm C were moved to Arm A if they failed to meet the early treatment completion criteria after bacteriological and radiological evaluation^13^. We performed longitudinal immune profiling on whole blood samples collected at baseline (week 0), during and at the end of the treatment (weeks 16 and 24), and one-year post-treatment (week 72). Whole blood was stimulated in TruCulture under four conditions: unstimulated control (Null), *M. tuberculosis*-antigen (*Mtb*-Ag; ESAT-6, CFP-10, and TB7.7 from QuantiFERON-TB Gold), lipopolysaccharide (LPS), and interleukin-1β (IL-1β). 25 consistently detectable proteins (cytokines, chemokines, growth factors) measured by Luminex multi- analyte profiling and low-abundance cytokines (IFN-γ, IL-2, pan-IFN-α, and IFN-β) by ultra- sensitive single-molecule array (Simoa) digital ELISA. Unless otherwise stated, all analyses focused on 33 patients with successful treatment outcome (Table 1).

**Table 1.** Patient characteristics.

| TB SATVI_Drug susceptible pulmonary TB patient characteristics |  |  |  |  |
| --- | --- | --- | --- | --- |
|  | Total | Arm A | Arm B | Arm C |
| Visit 0 (day) | 40 | 16 | 12 | 12 |
| Visit 16 (week) | 40 | 16 | 12 | 12 |
| Visit 24 (week) | 40 | 16 | 12 | 12 |
| Visit 72 (week) | 35 | 16 | 12 | 7 |
| Treatment successful | 33 (11F +22M) | 15 | 12 | 6 |
| Treatment failure | 2 (1F +1M) | 0 | 0 | 2 |
| Confirmed relapse | 4 (1F + 3M) | 0 | 0 | 4 |
| Reinfection | 1 (M) | 1 | 0 | 0 |
| Age, median (range), y | 37 (19-62) | 36 (19-49) | 40 (19-62) | 40 (26-57) |
| Sex | 13F + 27M | 5F + 11M | 5F + 7M | 3F + 9M |
| BMI at V0, median (range) | 18.0 (13.6-28.1) | 17.5 (14.7-23.0) | 18.3 (13.6-28.1) | 18.0 (15.8-21.4) |
| 2 previous TB episodes | 2 (M) | 1 | 0 | 1 |
| 1 previous TB episode | 14 (5F + 9M) | 4 | 6 | 4 |
| 0 previous TB episode | 24 (8F + 16M) | 11 | 6 | 7 |
| Smoker | 31 | 11 | 10 | 10 |
| Ex-smoker | 3 | 2 | 0 | 1 |
| Nonsmoker | 6 | 3 | 2 | 1 |
| **Exclusion criteria:** extrapulmonary TB, HIV-positive, COVID-positive, INH resistant, RIF resistance, pregnancy, diabetes, chronic inflammatory condition, use of immunosuppressive medications, etc.<br>F: Female; M: Male. |  |  |  |  |

### TruCulture *Mtb*-Ag stimulation reveals distinct protein responses in TB patients before, during and after antibiotic therapy

Principal component analysis (PCA) of 25 consistently detectable proteins revealed progressive convergence between Null and *Mtb*-Ag-stimulated protein profiles as patients recovered from TB disease (Figure 1A). We quantified this convergence by calculating centroid (Euclidean) distances between group centroids in PCA space (Fig. S2A). *Mtb*-Ag-stimulated profiles showed striking convergence toward Null, with centroid distance decreasing from 3.94 at baseline to 2.85 at treatment completion (Week 24; 28% reduction from baseline) and further to 1.50 by one-year post-treatment follow-up (Week 72; 62% reduction from baseline). IL-1β stimulation showed modest convergence (distance reduced from 2.39 at baseline to 2.26 at week 72; 5% reduction), while LPS-stimulated profiles remained consistently distinct throughout the study period (distance 8.51 at baseline to 8.87 at Week 72; 4% increase).

**Figure 1.**
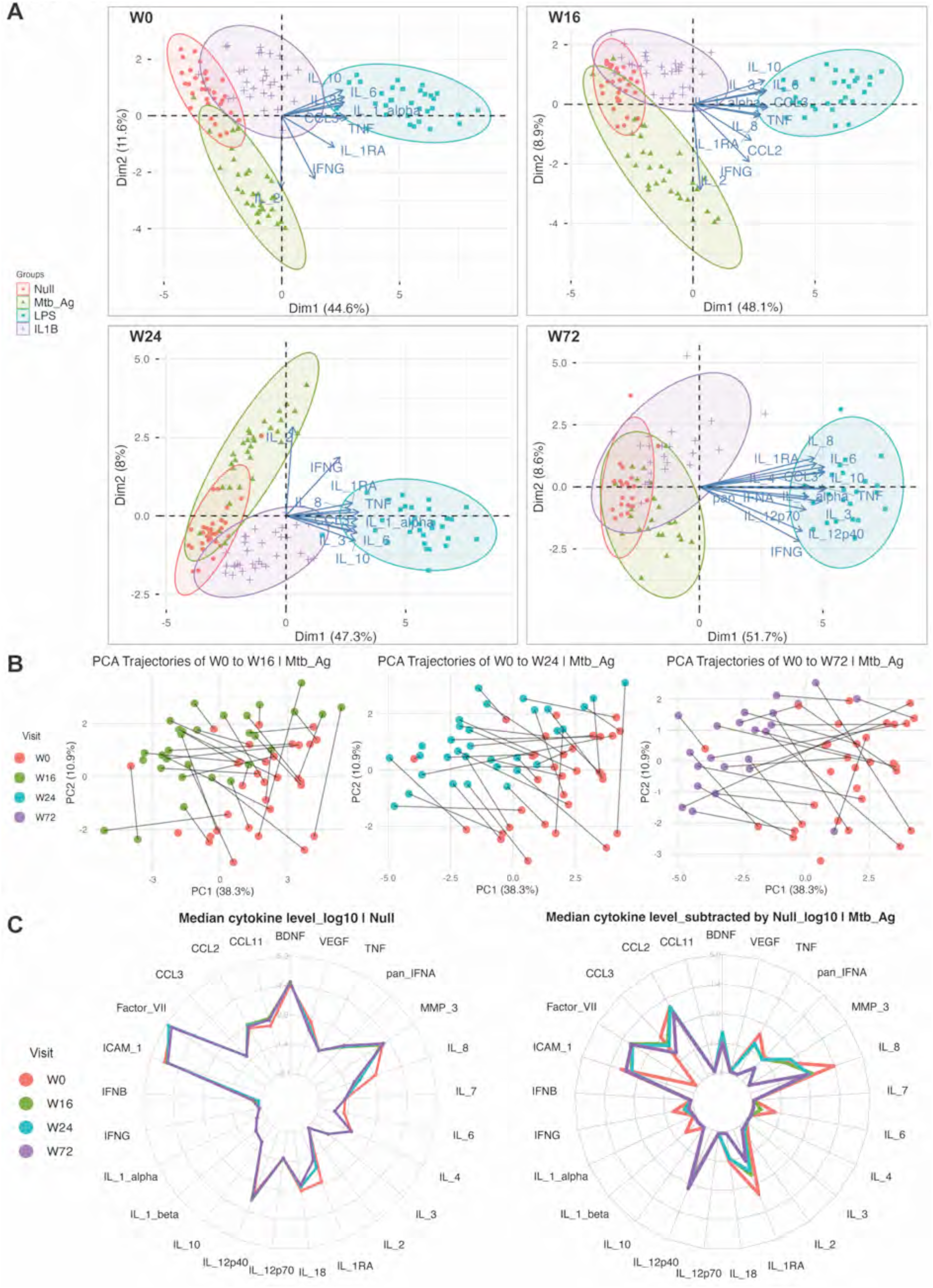
Luminex and Simoa cytokine profiles of TB patients before, during, and after successful antibiotic therapy. (A) Principal component analysis (PCA) of cytokine profiles under four TruCulture conditions (Null, *Mtb*-Ag, LPS, and IL-1β) at baseline (week 0), during treatment (weeks 16 and 24), and one-year follow-up post-treatment (week 72). Cytokines with cos2 > 0.7 are represented on the PCA plots. (B) Longitudinal PCA trajectories showing changes in cytokine profiles within individual patients over time, plotted using the same principal components for comparison. (C) Radar plots of median cytokine levels across visits. Cytokine levels in *Mtb*-Ag stimulation are the net values (stimulated subtracted by Null condition). Units from measurement: pan-IFN-α, fg/mL; all other cytokines, pg/mL.

To track individual patient responses, we performed PCA time-series analysis of longitudinal protein profiles. *Mtb*-Ag stimulation demonstrated the strongest treatment-associated differentiation among all conditions (Figure 1B). Protein profiles of individual patients progressively separated from baseline (week 0) during recovery, with 100% of week 72 profiles consistently separated from their respective baseline profiles in the same direction (Figure 1B). For individual cytokine levels, *Mtb*-Ag response among the 4 stimulations demonstrated the best discrimination of treatment (Figure 1C). This highlights the importance of using stimulation to amplify immune signals that remain undetectable or indistinguishable in conventional unstimulated patient samples.

### TruCulture stimulation identifies multiple stimulus-cytokine combinations that effectively reflect TB treatment response

To identify sensitive and reliable stimulus-cytokine combinations that distinguish between the pre-treatment timepoint and on-treatment timepoints at week 16 or 24, or the post- treatment timepoint at week 72, we applied Random Forest machine learning for feature importance ranking (Fig. S3A) and evaluated biomarker performance using area under the receiver operating characteristic curve (AUC-ROC). The top-ranked stimulus-cytokine combinations in Random Forest analysis also achieved the highest AUC-ROC scores (Figure 2A).

**Figure 2.**
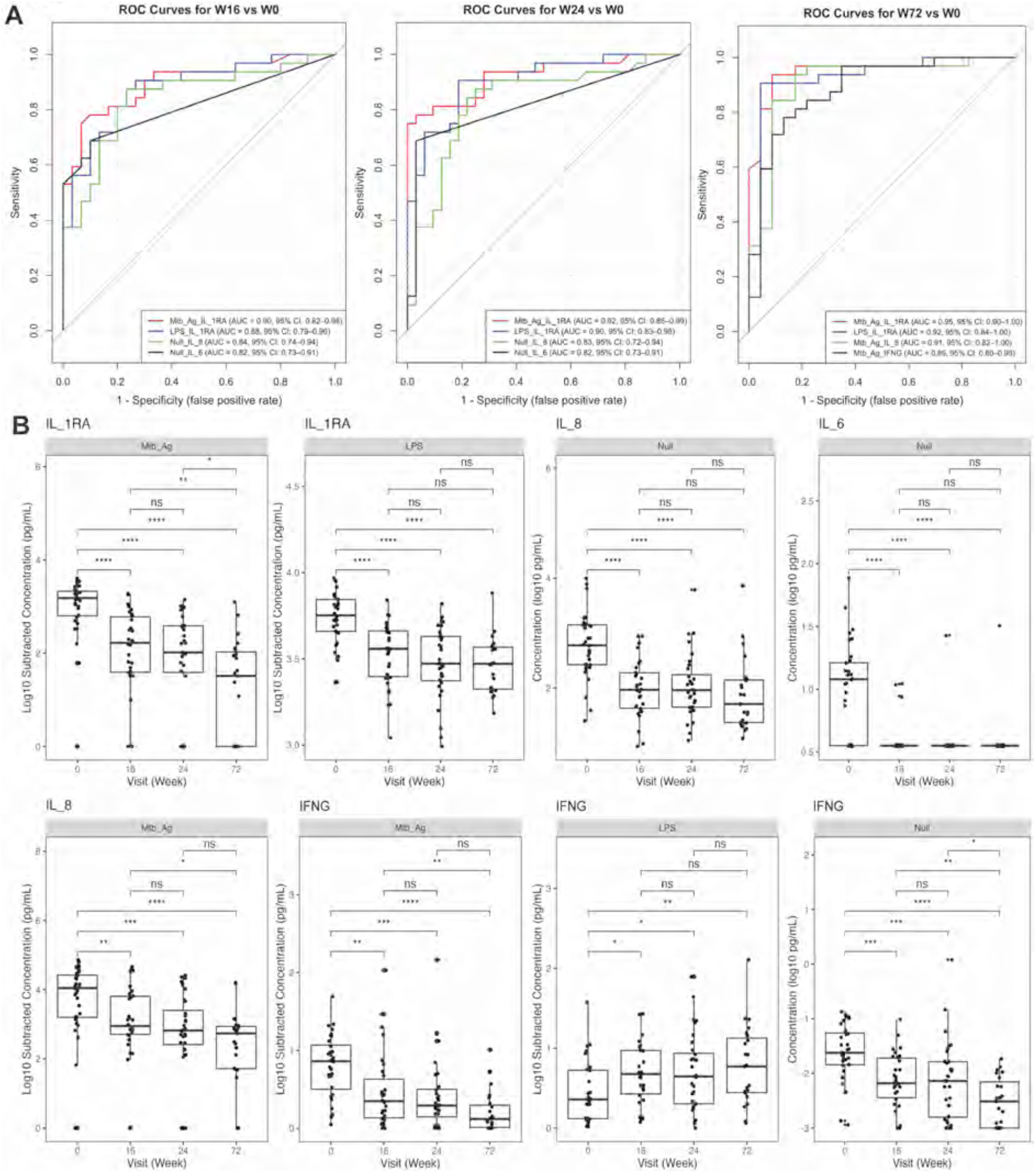
Cytokines for TB treatment monitoring. (A) Area under the receiver operating characteristic curve (AUC-ROC) for the top four stimulus–cytokine combinations identified by random forest machine learning model to distinguish untreated TB patients (W0) from treated patients (W16, W24, and W72). (B) Box plots of cytokines identified from random forest and AUC-ROC selection. Cytokine levels in *Mtb*-Ag and LPS stimulations show values subtracted by Null condition. IFNG concentration was measured by Simoa. P values were calculated using pairwise Wilcoxon tests with FDR adjustment. ns, not significant; *P < 0.05; **P < 0.01; ***P < 0.001; ****P < 0.0001.

*Mtb*-Ag-induced IL-1RA was the most powerful discriminatory protein biomarker, with AUC = 0.90 (95% CI: 0.82-0.98) at week 16, AUC = 0.92 (95% CI: 0.85-0.99) at week 24, and AUC = 0.95 (95% CI: 0.90-1.00) at week 72 (Figure 2A-B). Other biomarkers also demonstrated strong discrimination, including LPS-induced IL-1RA, Null-induced IL-8 and IL-6, and *Mtb*-Ag-induced IL-8 (Figure 2A-B). In an independent pulmonary TB cohort from Paris (MoSAR), Null IL-1RA and IL-18 also showed strong reductions following successful antibiotic treatment (Fig. S4A).

*Mtb*-Ag-induced IFN-γ, essential for host defense against *M. tuberculosis* and widely used to diagnose active TB disease^21,25^, achieved good discrimination at week 72 (AUC = 0.89; 95% CI: 0.80-0.98) but showed reduced performance at earlier timepoints (AUC = 0.73; 95% CI: 0.60- 0.86 at week 16. AUC = 0.78; 95% CI: 0.66-0.90 at week 24) (Figure 2A-B). While IFN-γ levels significantly reduced under Null and *Mtb*-Ag stimulation during TB recovery, LPS-induced IFN- γ significantly increased over time (Figure 2B). These opposing trends of decreased Null-IFN-γ and increased LPS-IFN-γ following treatment were also observed in our validation MoSAR TB cohort (Fig. S4A). This likely reflected distinct IFN-γ-secreting cell populations that respond differently to *Mtb*-Ag stimulation (CD4^+^, CD8^+^ T cells, and bystander activation by unconventional T cells)^26,27^ versus LPS stimulation (NK cells and monocytes)^28,29^.

Overall, 20 of 25 detectable cytokines changed significantly during treatment, with some continuing to evolve through post-treatment follow-up (Fig. S4B-E). Notably, all 11 cytokines previously identified as differentially expressed in active TB under *Mtb*-Ag stimulation^21^ exhibited consistent patterns in this study (Fig. S4C), confirming the reproducibility of the TruCulture whole blood stimulation system in TB disease studies. Intriguingly, despite antibiotic treatment ending at week 24 in Arm A and B, many unstimulated cytokines showed dramatic changes between week 0 and week 16, followed by less pronounced changes from week 16 to 72 (Fig. S5A). This kinetic pattern parallels sputum bacterial burden clearance, with no *M. tuberculosis* detected by Mycobacteria Growth Indicator Tube (MGIT) from week 16 onward in the treatment success group (Fig. S5B-C). The dramatic change of cytokine responses from week 0 to week 16 could be associated with *M. tuberculosis*-induced cell death and granuloma breakdown.

Despite the limited sample sizes in the treatment failure (n=2) and confirmed relapse (n=4) groups, we observed distinct cytokine response trajectories between different treatment outcome groups (Fig. S6A-B). Interestingly, all cytokine differences between treatment success and treatment failure groups occurred under *Mtb*-Ag stimulation (Fig. S6A). In contrast, most differences between treatment success and relapse groups occurred under IL-1β stimulation (Fig. S6B). This indicates that relapse is likely associated with innate immunity, particularly macrophage intracellular *M. tuberculosis* control^30^.

### Transcriptomic profiling identifies gene expression biomarkers of TB treatment response

To characterize transcriptional changes during TB treatment, we measured 815 immune genes using the NanoString Host Response Panel. Similar to the cytokine profiles, gene expression patterns under unstimulated (Null) conditions progressively converged with *Mtb*-Ag- stimulated profiles following treatment (Fig. S7A). Among all differentially expressed immune genes associated with TB treatment response (Fig. S7B), we screened stimulus_gene expression combinations by Random Forest machine learning (Fig. S3B) and validated their performance using AUC-ROC. The top-ranked stimulus-gene expression combinations in Random Forest analysis also achieved the highest AUC-ROC scores (Figure 3A). LPS-induced *NLRP3* gene expression was the most powerful discriminatory biomarker, with AUC = 0.94 (95% CI: 0.89 -1.00) at week 16 and good consistency across individuals (Figure 3B). NLRP3 is a critical component in inflammasome activation and pyroptosis, which plays an important role in *M. tuberculosis* inflammatory responses^31^. It has been reported that *M. tuberculosis*- induced plasma membrane damage causes NLRP3 activation and pyroptosis^32^. Although it is not yet clear whether inflammasome activation and pyroptosis are beneficial or detrimental to host defense against *M. tuberculosis*^31,33^.

**Figure 3.**
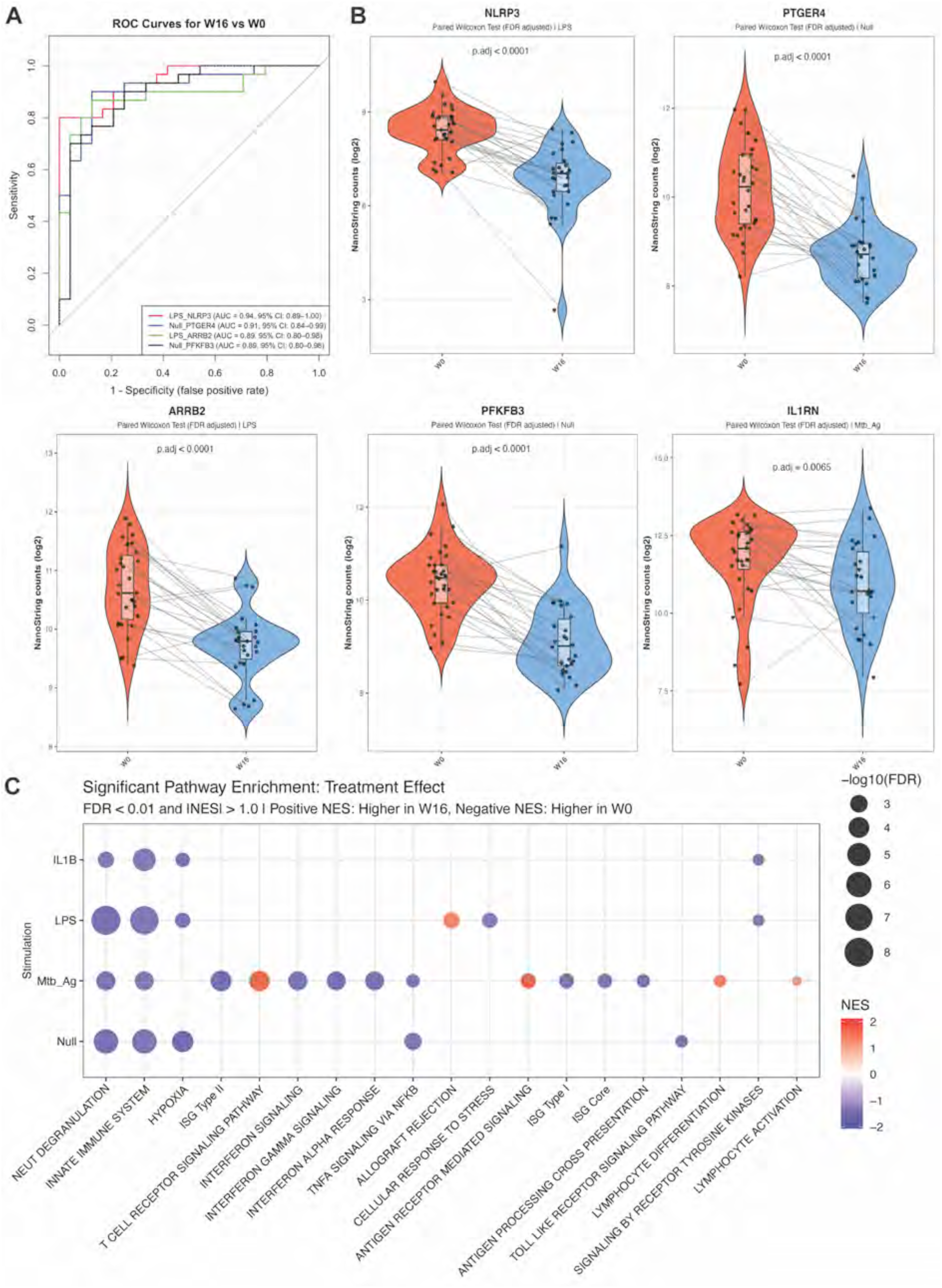
Immune gene expression in untreated and treated TB patients. (A) AUC-ROC plot and (B) violin plots for the top stimulus–gene expression combinations identified by random forest machine learning model to distinguish untreated patients (W0) from treated (W16) TB patients. (C) Bubble plot of gene set enrichment analysis showing gene signatures significantly altered between untreated (W0) and treated (W16) TB patients. Bubble color represents the normalized enrichment score (NES), and bubble size represents FDR-adjusted P values. Pathways are ordered by significance.

Other biomarkers including *PTGER4* (AUC = 0.91; 95% CI: 0.84 - 0.99), LPS-induced *ARRB2* (AUC = 0.89; 95% CI: 0.80 - 0.98), and *PFKFB3* (AUC = 0.89; 95% CI: 0.80 - 0.98] also exhibited good discriminatory power (Figure 3B) for TB treatment response. Consistent with the protein-level IL-1RA results, *Mtb*-Ag-induced *IL1RN* gene expression also showed significant (Wilcoxon FDR p=0.0065) distinction for pre-treatment and on-treatment timepoints (Figure 3B; AUC = 0.68; 95% CI: 0.53-0.83).

To identify changes in biological pathways during treatment response, we performed gene set enrichment analysis (GSEA) on differentially expressed genes (DEGs) between pre-treatment (W0) and on-treatment (W16) TB patients. Following treatment, active TB patients exhibited significantly decreased expression of inflammatory signatures (innate immune system, neutrophil degranulation, hypoxia, IFN-γ signaling, IFN-α response, ISG signaling, TNF-α-NF- κB signaling, cellular stress response, antigen processing-cross presentation, signaling by receptor tyrosine kinases, toll-like receptor signaling (Figure 3C, Table S1). Conversely, lymphocyte signatures (T cell receptor signaling, antigen receptor-mediated signaling, lymphocyte differentiation and activation) were significantly elevated in active TB patients following treatment (Figure 3C, Table S1).

### Integration of FDG PET/CT scan lung lesions with blood biomarkers

FDG PET/CT imaging was used in the Predict-TB trial to stratify patients by disease severity and predict treatment outcomes^13^. Both total cavity volume and total lesion glycolysis significantly decreased during treatment (Figure 4A). While total cavity and total lesion glycolysis were significantly correlated, total lesion glycolysis remained detectable and variable even in patients without lung cavities (Fig. S8A).

**Figure 4.**
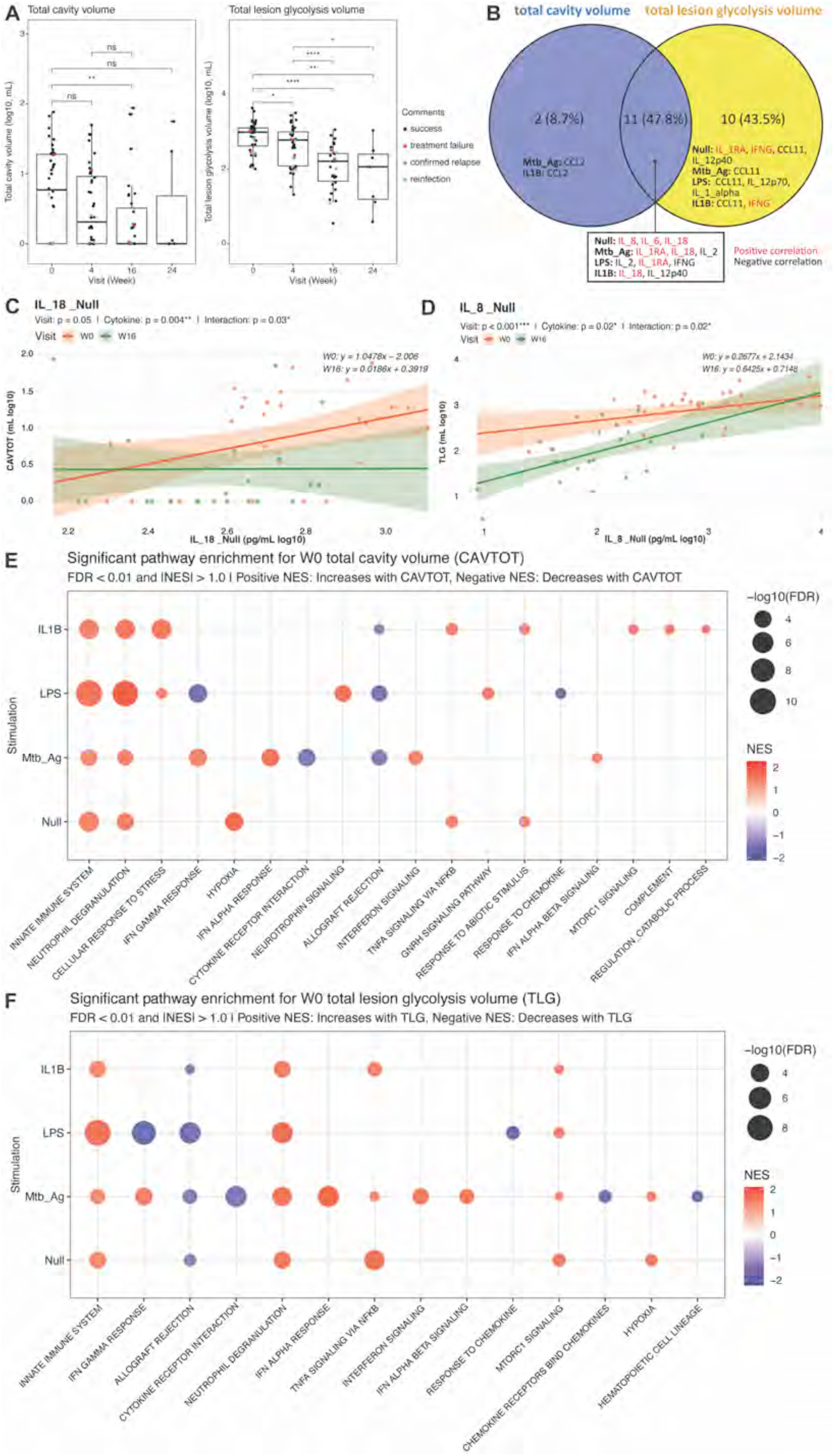
Integration of FDG PET/CT scan results with blood immune responses. (A) Box plots showing changes in lung lesion volume as TB patients recovered from TB. P values were calculated using pairwise Wilcoxon tests with FDR adjustment. ns, not significant; *P < 0.05; **P < 0.01; ***P < 0.001; ****P < 0.0001. (B) Venn diagram of W0 stimulus–cytokine combinations significantly associated with W0 TB lung lesions, validated by at least two models (random forest, linear model, and Spearman correlation). (C) Linear mixed-effects models of significant cytokine and treatment effects on total cavity volume (CAVTOT), with 95% confidence interval. P values were FDR-adjusted. (D) Linear mixed-effects models of significant cytokine and treatment effects on total lesion glycolysis volume (TLG), with 95% confidence interval. P values were FDR-adjusted. Bubble plot of gene set enrichment analysis showing gene signatures significantly associated with (E) total cavity volume and (F) total lesion glycolysis volume, in untreated TB patients. Bubble color represents the normalized enrichment score (NES), and bubble size represents FDR-adjusted P values. Pathways are ordered by significance.

To identify systemic immune biomarkers associated with the extent of lung pathology in untreated (W0) TB patients, we applied Random Forest analysis to screen for cytokine- stimulus combinations correlated with total cavity and total lesion glycolysis volumes (Fig. S8B). Candidates were validated using linear and Spearman correlation models (Fig. S8C-D). Biomarkers identified by at least two models were considered high-confidence associations (Figure 4B). Previously identified treatment-responsive biomarkers, including Null-induced IL- 8, IL-6, IL-18, *Mtb*-Ag-induced IL-1RA and LPS-induced IL-1RA, exhibited positive correlations with both total cavity volume and total lesion glycolysis volume (Figure 4B). CCL2 (MCP-1) negatively correlated with total cavity volume under *Mtb*-Ag and IL-1β stimulations (Figure 4B). IFN-γ showed positive correlation with total lesion glycolysis under Null and IL-1β stimulation, while CCL11 (eotaxin-1) exhibited negative correlation with lesion glycolysis across all four stimulation conditions (Figure 4B).

To investigate cytokine-treatment effects on TB lung lesions, we constructed a linear mixed-effects (LME) model to leverage the multiple measurements available per patient. IL-18 showed significant positive correlation with total cavity volume at the pre-treatment (W0) timepoint but not at the on-treatment (W16) timepoint (Figure 4C). In contrast, IL-8 showed a stronger positive correlation with total lesion glycolysis volume in at W16 than at the pre-treatment (W0) timepoint (Figure 4D).

To identify biological pathways associated with TB lung lesions, we performed GSEA on gene expression correlated with lung cavity and total lesion glycolysis at the pre-treatment (W0) timepoint. Total cavity volume was associated with elevated inflammatory signatures (innate immune system, neutrophil degranulation, cellular response to stress, *Mtb*-Ag-induced IFN-γ response, hypoxia, IFN-α response, neurotrophin signaling, TNF-α-NF-κB signaling, GnRH signaling, response to abiotic stimulus, IFN-α/β signaling, mTORC1 signaling, complement, regulation of catabolic process), and reduced signatures in LPS-induced IFN-γ response, cytokine-cytokine receptor interaction, allograft rejection, and response to chemokine (Figure 4E, Table S1). Total lesion glycolysis showed similar patterns, with elevated inflammatory signatures (innate immune system, neutrophil degranulation, IFN-α response, TNF-α-NF-κB signaling, *Mtb*-Ag-induced IFN-γ response, IFN-α/β signaling, mTORC1 signaling, hypoxia), alongside reduced signatures in LPS-induced IFN-γ response, allograft rejection, cytokine- cytokine receptor interaction, response to chemokine, chemokine receptors bind chemokines, and hematopoietic cell lineage (Figure 4F, Table S1).

Week 16 total lung cavity volume and lesion glycolysis volume were recently found to predict unfavorable outcomes in a larger group of Predict-TB patients^13^. To identify biological pathways associated with lung lesion-mediated unfavorable outcomes, we performed GSEA on gene expression correlated with week 16 lung cavity and total lesion glycolysis. Week 16 total cavity volume was associated with elevated innate immune signatures (neutrophil degranulation, mTORC1 signaling, IFN-α response, hypoxia), and decreased inflammatory signatures (TNF-α-NF-κB signaling, cytokine-cytokine receptor interaction, allograft rejection, NLR signaling, regulated necrosis) (Fig. S9A). Week 16 total lesion glycolysis correlated with elevated inflammatory pathways (TNF-α-NF-κB, IFN-γ response, IFN-α response, hypoxia, neutrophil degranulation), and reduced signature in intracellular signaling by second messengers (Fig. S9B, Table S1).

### Demographic and clinical factors influence TB disease and treatment response

Older age, male sex, low BMI, a history of previous TB, and higher bacterial burden are known TB risk factors ^34–36^. We evaluated their impact on lung pathology and systemic immune biomarkers. Multicollinearity among tested demographic and clinical factors was minimal (Fig. S10A). For the impact of age, linear mixed-effects modeling revealed that both total cavity volume and total lesion glycolysis decreased significantly following treatment in younger but not older patients (Figure 5A). Similarly, LPS-induced IFN-γ increased significantly with treatment only in younger patients (Figure 5B). Expression of several immune genes (*BNIP3, CXCR4, DUSP3, IL2RG, MIF, PGK1, PIK3CB, RBPJ*) also showed treatment-associated decreases exclusively in younger patients (Fig. S10B). Intriguingly, these age-treatment interactions were observed only under IL-1β stimulation (Fig. S10B), suggesting age-dependent effects on innate immunity. Comparison of GSEA between untreated (W0) and treated (W16) individuals identified TB-specific pathways correlated with age (Figure 5C and Fig. S11A). In untreated TB patients, age positively correlated with gene expression signatures in asthma, lymphocyte chemotaxis, and response to chemokine, while negatively correlated with signatures in cellular response to stress, SARS-COV-2 infections and host interactions (Figure 5C, Table S1). These age-associated gene expression patterns were attenuated or absent following treatment (Fig. S11A, Table S1).

**Figure 5.**
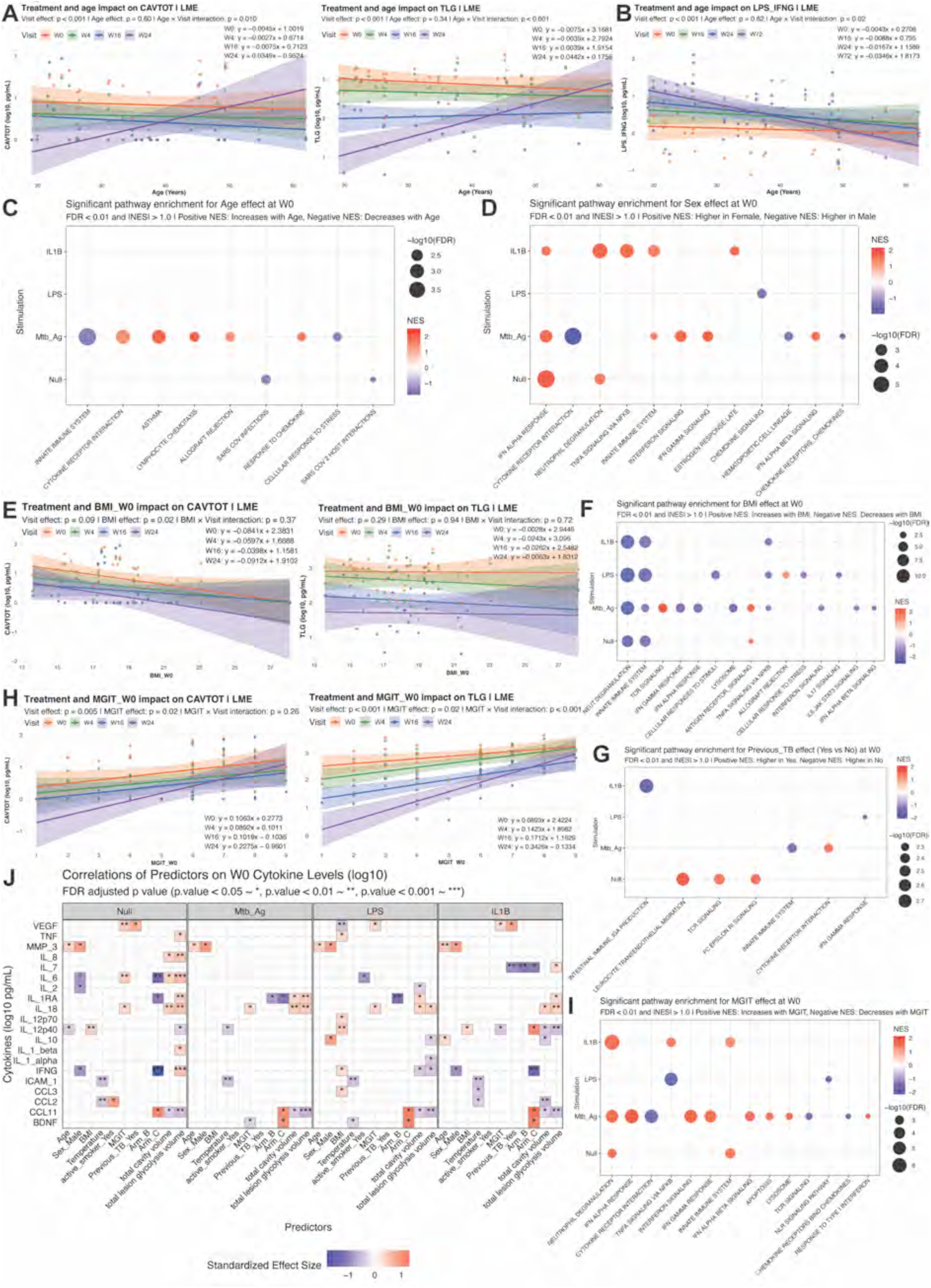
Impact of demographic and clinical factors on TB patients. (A) Linear mixed-effects models of age and treatment effects on total cavity volume (CAVTOT) and total lesion glycolysis volume (TLG), with 95% confidence interval. (B) Linear mixed-effects models of significant age and treatment effects on IFNG level under LPS condition, with 95% confidence interval. (C) Bubble plot of gene set enrichment analysis showing gene signatures significantly associated with age in W0 untreated TB patients. Pathways are ordered by significance. (D) Bubble plot of gene set enrichment analysis showing gene signatures significantly associated with sex in W0 untreated TB patients. Pathways are ordered by significance. (E) Linear mixed- effects models of BMI and treatment effects on total cavity volume (CAVTOT) and total lesion glycolysis volume (TLG), with 95% confidence interval. (F) Bubble plot of gene set enrichment analysis showing gene signatures significantly associated with BMI in W0 untreated TB patients. Pathways are ordered by significance. (G) Bubble plot of gene set enrichment analysis showing gene signatures significantly altered between W0 untreated TB patients with and without a history of at least one previous TB episode more than three years before enrollment. Pathways are ordered by significance. (H) Linear mixed-effects models of *M. tuberculosis* burden (MGIT) and treatment effects on total cavity volume (CAVTOT) and total lesion glycolysis volume (TLG), with 95% confidence interval. (I) Bubble plot of gene set enrichment analysis showing gene signatures significantly associated with *M. tuberculosis* burden in W0 untreated TB patients. Pathways are ordered by significance. (J) Linear models identifying demographic and clinical factors significantly associated with W0 cytokine levels across different stimulation conditions.

To investigate the impact of biological sex on TB disease, we evaluated sex effects in lung lesion and blood biomarkers. Lung lesion severity did not differ significantly between male and female patients (Fig. S10C). Random Forest and limma DEG analysis identified TB-specific sex differences in *CCR4* gene expression under Null, IL-1β and *Mtb*-Ag stimulations (Fig. S10D). Comparison of GSEA between untreated (W0) and treated (W16) individuals identified TB- specific pathways correlated with sex (Figure 5D and Fig. S11B). Female TB patients had elevated signatures in type 1 interferon response (IFN-α, IFN-β signaling) and estrogen response, while male TB patients had elevated gene expression signatures in chemokine- receptor interaction, chemokine signaling, and hematopoietic cell lineage (Figure 5D, Table S1). These sex-associated gene expression patterns were attenuated or absent following treatment (Fig. S11B, Table S1).

BMI increased progressively during successful treatment (Fig. S10E) without sex differences (Fig. S10F). Linear mixed-effect models showed that in untreated (W0) TB patients, BMI negatively correlated with total cavity volume but not total lesion glycolysis (Figure 5E). Comparison of GSEA between pre-treatment (W0) and on-treatment (W16) timepoints identified TB-specific pathways correlated with BMI (Figure 5F and Fig. S11C). Pre-treatment, BMI positively correlated with gene expression signatures in T cell receptor signaling, antigen receptor signaling, and allograft rejection, while negatively correlated with signatures in cellular response to stimuli, lysosome, TNF-α-NF-κB signaling, vesicle-mediated transport, cellular response to stress, IL-17 signaling, and IL-6-JAK-STAT3 signaling (Figure 5F, Table S1). These BMI-associated gene expression patterns were attenuated or absent following treatment (Fig. S11C, Table S1).

Previous TB history did not impact extent of lung lesions (Fig. S10G). Comparison of GSEA between pre-treatment (W0) and on-treatment (W16) timepoints identified TB-specific pathways correlated with presence of previous TB (Figure 5G and Fig. S11D). Pre-treatment, previous TB positively correlated with gene expression signatures in leukocyte transendothelial migration and Fc epsilon RI-mediated signaling, while negatively correlated with intestinal immune network for IgA production (Figure 5G, Table S1). These previous TB - associated gene expression patterns were attenuated or absent following treatment (Fig. S11D, Table S1).

Sputum *M. tuberculosis* burden became undetectable from week 16 following successful treatment (Fig. S5C). Both cavity volume and lesion glycolysis volume correlated positively with sputum *M. tuberculosis* burden (MGIT) (Figure 5H). At the pre-treatment timepoint, GSEA showed that bacterial burden positively associated with inflammatory signatures (innate immune system, neutrophil degranulation, IFN-γ signaling, IFN-α signaling, IFN-β signaling), apoptosis, and lysosome signaling, while negatively associated with cytokine-cytokine receptor interaction, T cell receptor signaling, NLR signaling, chemokine receptors bind chemokines (Figure 5I, Table S1).

Linear modeling identified multiple stimulus-cytokine combinations associated with demographic factors (age, sex, BMI, body temperature, smoking status) and clinical parameters (*M. tuberculosis* burden, previous TB, disease severity, lung lesions) at pre-treatment (W0) (Figure 5J). Some associations, including elevated MMP3 in males and negative correlation of CCL11 with disease severity, persisted at week 16, while most associations were specific to the pre-treatment timepoint (Figure 5J, Fig. S10H).

## Discussion

Our study demonstrates that TruCulture whole blood stimulation is a powerful platform for TB immunomonitoring and provides insights for improved patient stratification. We identified systemic blood biomarkers significantly associated with TB treatment responses, lung lesion extent, and demographic and clinical factors. These findings have important implications for personalized TB care through early prediction of treatment responses and real-time therapy monitoring. Furthermore, the gene expression signatures we identified reveal pathways involved in TB pathology and patient heterogeneity.

Older age, male sex, and higher bacterial burden emerged as the strongest demographic and clinical factors influencing treatment responses in this study (Figure 5, Fig. S10). The impact of bacterial burden on TB disease progression and treatment responses is well-established in both animal models and clinical studies^37,38^. Similarly, aging is a recognized risk factor across many diseases, broadly affecting vaccine and treatment efficacy^39,40^. The male bias in TB susceptibility, however, requires particular attention given accumulating evidence for a biological basis beyond socioeconomic and behavioral factors. TB affects males disproportionately (male-to-female incidence ratio 1.7) with greater severity and mortality^36^. Several lines of evidence suggest biological sex plays a critical role in TB: females exhibit stronger BCG immune reactivity^36^, male mice are more susceptible to aerosol *Mtb* infection (reduced survival, higher bacterial burden, worse clinical symptoms)^41^, and castrated males have lower TB mortality (6/ 74, 8.1%) compared to gonadal males (35/170, 20.6%)^36^. Linear modeling revealed that male patients had significantly lower IFN-γ and higher LPS-induced IL- 10 levels than female patients at the pre-treatment timepoint (Figure 5J). This is consistent with findings in murine *Mtb* aerosol infection model^42^ and testosterone’s known suppression of IFN-γ and enhancement of IL-10 production^36^.

Among all sites in the Predict-TB study^13,24^, the SATVI site showed a disproportionately high rate of unfavorable outcomes compared to the other four Cape Town sites. Although SATVI accounted for approximately 16% of total enrollment, it contributed 36% of all unfavorable outcome cases. Especially in Arm C, SATVI accounted for 50% of unfavorable outcomes across South African sites. Exploratory analysis did not identify significant differences in common risk factors, medical history, or baseline clinical features between sites. Some unfavorable outcome cases from SATVI in Arm C even had low or minimal baseline PET/CT readouts. A key demographic distinction between SATVI and other Cape Town sites is the rural versus urban setting. The higher rate of unfavorable outcomes at SATVI may reflect the impact of socioeconomic disadvantages associated with rural area, as socioeconomic status can impact immune responses in humans^43^. Ongoing whole-genome sequencing (WGS) of *Mtb* strains from rural SATVI and urban Cape Town cohorts may provide additional insights into whether strain-specific factors contribute to the increased unfavorable outcome in SATVI.

There is a growing awareness that immune reconstitution continues beyond pathogen clearance^44,45^. Post-treatment complications, in particular persistent lung impairment, affect up to 75% of successfully treated TB patients^46^. Previous studies of paired blood and bronchoalveolar lavage fluid (BALF) identified a few proinflammatory cytokines correlated with post-TB lung disease^46^. We observed continued evolution of multiple stimulus-cytokine combinations during the post-treatment period (weeks 24-72), including Null-induced IFN-γ, *Mtb*-Ag-induced IL-1RA, TNF-α, CCL2, CCL3, and LPS-induced IL-3 (Figure 2B, Fig. S4C-D). These findings extend previous observations and suggest that prolonged immune monitoring can improve understanding of post-TB treatment complications and enhance patient care strategies.

We identified *Mtb*-Ag-induced IL-1RA as the best protein biomarker that reflected successful treatment response. This finding is consistent with previous reports of elevated plasma IL-1RA levels and dysregulated IL-1 responses in active TB patients compared to asymptomatic QFT- positive controls^20^. IL-1RA functions as a competitive inhibitor of IL-1α and IL-1β by binding to IL-1R1, thereby counterbalancing IL-1-mediated host protection^47^. Studies in mice demonstrated that type I IFN-driven susceptibility to *M. tuberculosis* infection is mediated through IL-1RA^48^, highlighting its dual role as both a biomarker and immunoregulatory mediator in TB pathogenesis. In addition to *Mtb*-Ag-induced IL-1RA, we also identified LPS- induced *NLRP3* as the best transcriptomic biomarker, and Null-IL-18 as a strong protein biomarker (validated in both SATVI and MoSAR cohorts) for distinguishing treated from untreated patients. Together, these findings strongly implicate the involvement of inflammasome activation and pyroptosis in TB disease.

Gene expression analysis revealed multiple biological pathways associated with TB treatment response and disease pathology. Notably, several identified pathways have context-dependent effects in TB pathogenesis. Neutrophils contribute to early pathogen control through oxidative killing of phagocytosed mycobacteria^49^ but drive immunopathology and correlate with disease severity when excessive^50^. Similarly, TNF confers resistance to intracellular *M. tuberculosis* during early infection but causes detrimental cell death and tissue damage when excessive^51^. Pyroptosis follows a similar pattern: controlled pyroptotic signaling releases danger signals and amplifies protective immunity^33^, while excessive pyroptosis during late infection drives tissue damage and facilitates bacterial dissemination^33^. The significantly elevated lymphocyte signatures observed in active TB patients following treatment (Figure 3C) are likely due to restoration of T cell exhaustion and B cell dysfunction reported in TB disease^52,53^. Future mechanistic studies are required to decipher if the upregulation or downregulation of the identified pathways are beneficial or detrimental to TB pathogenesis.

The ability of whole blood stimulation to distinguish active TB^21^, track treatment responses, and stratify patients suggests its potential utility for TB vaccine evaluation, which lacks reliable and sensitive biomarkers^54^. Limitations of this study include small sample size and difficulty in identifying replication cohorts with comparable trial designs. Despite these limitations, our study establishes TruCulture whole blood stimulation as a valuable platform for TB immunomonitoring, identifies candidate biomarkers for treatment response and disease severity, and provides mechanistic insights into dynamic immune responses during TB therapy and personalized patient care.

## Materials and Methods

### Clinical study design

This study was nested within the Predict-TB clinical trial (clinicaltrials.gov NCT02821832) to identify biomarkers to predict TB treatment responses^13,24^. Briefly, adult patients with active pulmonary TB disease were enrolled at the South African Tuberculosis Vaccine Initiative (SATVI) trial site in Worcester. Inclusion and exclusion criteria were described in Table 1 and detailed previously^13,24^. Eligible participants initially received two months of the “intensive phase” treatment regimen with four drugs daily (isoniazid, rifampicin, pyrazinamide and ethambutol) followed by a two drug (isoniazid and rifampicin) continuation phase until treatment completion. Following baseline FDG PET/CT evaluation, TB patients were stratified by predicted relapse risk and assigned to three treatment arms. High-risk patients received 24- weeks of standard treatment (Arm A, n=16). Low-risk patients were randomized to either 24- week (Arm B, n=12) or 16-week (Arm C, n=12) regimens. At week 16, participants in Arm C were moved to Arm A if they failed to meet the early completion criteria after bacteriological and radiological evaluation^13^. Written, informed consent was obtained in the local language of the participant. The trial was reviewed and approved by the IRB/ethics committees of the NIAID, Stellenbosch University, Faculty of Health Sciences University of Cape Town, and South African Health Products Regulatory Agency.

For the repeatability studies asymptomatic QFT-positive donors (n=9) and individuals with bacteriologically confirmed active pulmonary TB (n=1) were recruited at the South African Tuberculosis Vaccine Initiative (SATVI) site or public health clinics in Worcester, and an additional 4 active TB disease patients were recruited at Cochin Hospital in Paris. All participants were HIV-negative. Blood was stimulated in 4 technical replicates to assess repeatability, and classification cut-offs were defined from a previous TruCulture TB study^21^. For reproducibility testing, asymptomatic QFT-positive donors (n=12) recruited at SATVI, were sampled at 4 independent timepoints each 4 weeks apart. Reproducibility testing was not performed on active TB disease patients. Two different TruCulture tube batches were also tested with no differences in patient classification observed due to this batch. Only Null and *Mtb*-Ag conditions were available in repeatability and reproducibility testing.

Active pulmonary TB patients in the validation MoSAR cohort (ClinicalTrials.gov ID NCT05916638) were recruited from the departments of internal medicine, pneumology, rheumatology, and infectious diseases at a tertiary referral university hospital (Hôpital Bichat, Assistance Publique–Hôpitaux Paris, Paris, France). Affiliation to medical insurance was required. Exclusion criteria for active pulmonary TB patients were HIV infection, chronic inflammatory condition (sarcoidosis), and pregnancy to avoid potential confounding of immunological phenotypes. Paired blood samples from 12 TB patients collected before treatment initiation and after 24 weeks of successful antibiotic therapy were used to validate biomarkers identified in the SATVI cohort. Only Null and LPS conditions were available in MoSAR TB treatment response cohort. Epidemiological, demographic, clinical, laboratory, imaging, and pathological data, along with treatment and follow-up information, were collected using a standardized REDCap electronic data capture form. Written informed consent was obtained from all participants.

### Treatment outcome determination

Treatment outcomes were defined based on bacteriological testing of sputum samples as previously described^24^. Briefly, treatment success was defined as at least two consecutive negative cultures (at least four weeks apart) before treatment completion without subsequent positivity. Treatment failure was defined as culture-positivity at week 24 or confirmed reconversion before week 24. Recurrence was culture reconversion after week 24, confirmed by repeat culture. Relapses (same strain) were distinguished from reinfections (different strain) by MIRU-VNTR genotyping and whole-genome sequencing. Only relapses were considered as study endpoints.

### Mycobacterium tuberculosis detection

GeneXpert MTB/RIF test was used to detect the presence of *M. tuberculosis* in sputum samples and to determine rifampin susceptibility. Solid culture (Löwenstein–Jensen medium) was used to determine treatment outcome. Time to detection (TTD) measurements from Mycobacteria Growth Indicator Tube (MGIT) liquid culture were available in all patients at all study time points and were therefore used as sputum *M. tuberculosis* burden indicator in this study. MGIT liquid culture was also used in clinical assessment at enrolment and confirmation of TB treatment outcome.

### TruCulture Whole-blood Stimulations

TruCulture tubes were prepared in batch following standardized procedures as previously described^16^. Briefly, TruCulture buffered media (2 ml) were resuspended with the following stimuli: Null, *M. tuberculosis* antigen (ESAT-6, CFP-10, and TB7.7 from QuantiFERON-TB Gold), LPS (Invivogen, 10 ng/ml), and IL-1β (Peprotec, 25 ng/ml). Prepared tubes were stored at −20 °C until use. Peripheral blood was collected in sodium-heparin tubes (50 IU/ml final concentration). Within 30 minutes of collection, 1 mL of whole blood was transferred into pre-warmed TruCulture tubes, placed in a dry block incubator, and incubated at 37 °C for 22 hours. Following incubation, a valve was inserted to separate the cellular and supernatant compartments to terminate stimulation reaction. Supernatants were removed for protein profiling, and cell pellets were resuspended in 2 mL of TRIzol LS reagent (Sigma) for transcriptomic analyses. Samples were vortexed for 2 minutes, allowed to stand for 10 minutes at room temperature, and subsequently stored at −80 °C until further processing.

### Multi-analyte protein profiling

Supernatants from TruCulture tubes were used for protein profiling. 25 consistently detectable proteins including cytokines, chemokines and growth factors were measured by Luminex xMAP technology (Rules-Based Medicine) as previously described^15,21^. Low- concentration cytokines (IFN-γ, IL-2, pan-IFN-α, and IFN-β) were measured by ultra-sensitive digital ELISA Simoa as previously described^55–57^.

### Nanostring Transcriptomics

Trizol stabilized TruCulture cell pellets were used for RNA extraction and NanoString gene expression analysis. RNA was quantified using the Qubit RNA HS Assay Kit (Thermo Fischer Scientific), and RNA integrity was assessed with Agilent 2200 TapeStation system. NanoString nCounter system was used for the digital counting of transcripts of 815 genes in NanoString nCounter Host Response Panel, including 30 customised genes. Quality control of the data was done by checking the following metrics using the nSolver^TM^ analysis software (NanoString technologies): fields of view counted (flag if < 0.75), binding density (flag if not in the 0.05- 2.75 range), linearity of positive controls (flag if R2 < 0.9) and limit of detection for positive controls (flag if 0.5 fM positive control < 2 SDs above the mean of the negative controls). Gene expression counts were normalized following background subtraction of 8 negative control probes, using 6 positive control probes and 6 housekeeping genes (MRPS7, NMT1, SDHA, GUSB, HPRT1, ABCF1) selected by the ‘geNorm’ algorithm in R and 100% detectability in all tested samples.

### Gene set enrichment analysis

Gene set enrichment analysis (GSEA) was performed on 815 genes from NanoString nCounter Host Response Panel to identify biological pathways associated with demographic and clinical variables. For each variable, gene expression association statistics were obtained using linear modelling in R (limma package). Continuous variables (age, BMI, MGIT, total cavity volume, and total lesion glycolysis volume) were modelled using linear regression of normalized gene expression on the variable. Binary variables (sex and previous TB) were modeled using two-group comparisons with the binary factor coded as an indicator variable. For the treatment status comparison (W0 vs W16), a paired analysis was performed accounting for within-donor correlation by including donor as a blocking factor in the linear model design matrix. For each variable, genes were ranked by the t-statistic corresponding to the variable’s regression coefficient, with positive values indicating higher expression associated with increasing variable values (or in the reference group for binary variables). Ranked gene lists were used as input for pre-ranked GSEA against curated gene sets from the Molecular Signatures Database (MSigDB, collection: Hallmark, KEGG, Reactome, GO Biological Processes). Pathways with false discovery rate (FDR) < 0.01 and normalized enrichment score (NES) absolute value > 1.0 were considered significantly enriched.

### FDG PET/CT imaging

Lung lesions of TB patients were measured by fludeoxyglucose-18 (FDG) positron emission tomography (PET) and computed tomography (CT) as previously described^13^. Briefly, participants fasted for at least 6 hours before [18F] FDG administration. According to body weight, 185–259 Megabecquerel (MBq) of [18F] FDG was administered intravenously 60 minutes before the scan. The PET/CT scan images were segmented using an enhanced version of automated computational method followed by manual inspection as previously described^13^.

### Statistical analysis

All statistical analyses were performed using R (version 4.4.2). Data normality was evaluated by Shapiro-Wilks test. Unless otherwise stated, all variables were analysed using non-parametric methods due to non-normal distributions in most datasets and modest sample sizes. Random Forest machine learning algorithm was used from the ’randomForest’ package in R (version 4.7-1.2). The model was trained for classification and regression of categorical and numerical variables. Model performance was assessed by 5-fold cross-validation with 3 repetitions. 10 random noise variables with normal distribution while matching the study dataset characteristics were generated by ‘rnorm’ function as a negative control. Variables with importance scores exceeding the average importance score across the random variables were considered as biologically meaningful predictors. Other main R packages used in this study were: ggplot2 (graphics), FactoMineR (PCA), lme4 (linear mixed-effects model), limma (differential gene expression analysis), and fgsea (gene set enrichment analysis). Multiple comparison corrections were performed using the Benjamini-Hochberg false discovery rate (FDR) method.

## Acknowledgments

The study participants and the SATVI, MoSAR, and Cochin clinical and laboratory teams.

UTechS CB of the Center for Translational Research, Institut Pasteur for supporting Nanostring experiments.

J.F. thanks E. Villain and A. Desrentes for bioinformatics discussions, Q. Richier and B. Jean for clinical TB discussions.

## Grant information

D.D. acknowledges support from the Gates Foundation grant [INV- 007585]. The Predict TB clinical study was supported by the Gates Foundation [OPP1155128]; and the European and Developing Countries Clinical Trials Partnership [SRIA.2015.1065].

J.F. acknowledges support from Lady Mireille and Sir Dennis Gillings Global Public Health Fellowship. M.R. was supported by a grant from the Fondation pour la Recherche Médicale (FDM202206015344). She received “La Bourse Junior” from the Société de Pathologie Infectieuse de Langue Française (SPILF) and the “Bourse de Recherche” from Le Fonds de dotation Villa M, Groupe Pasteur Mutualité et ses mutuelles partenaires. M.R. is member of the Ecole de l’Inserm Liliane Bettencourt Programme.

## Conflicts of interest statement

D.D. has received grant support from Rules Based Medicine unrelated to this study.

## Supplemental Information

**Fig. S1.**
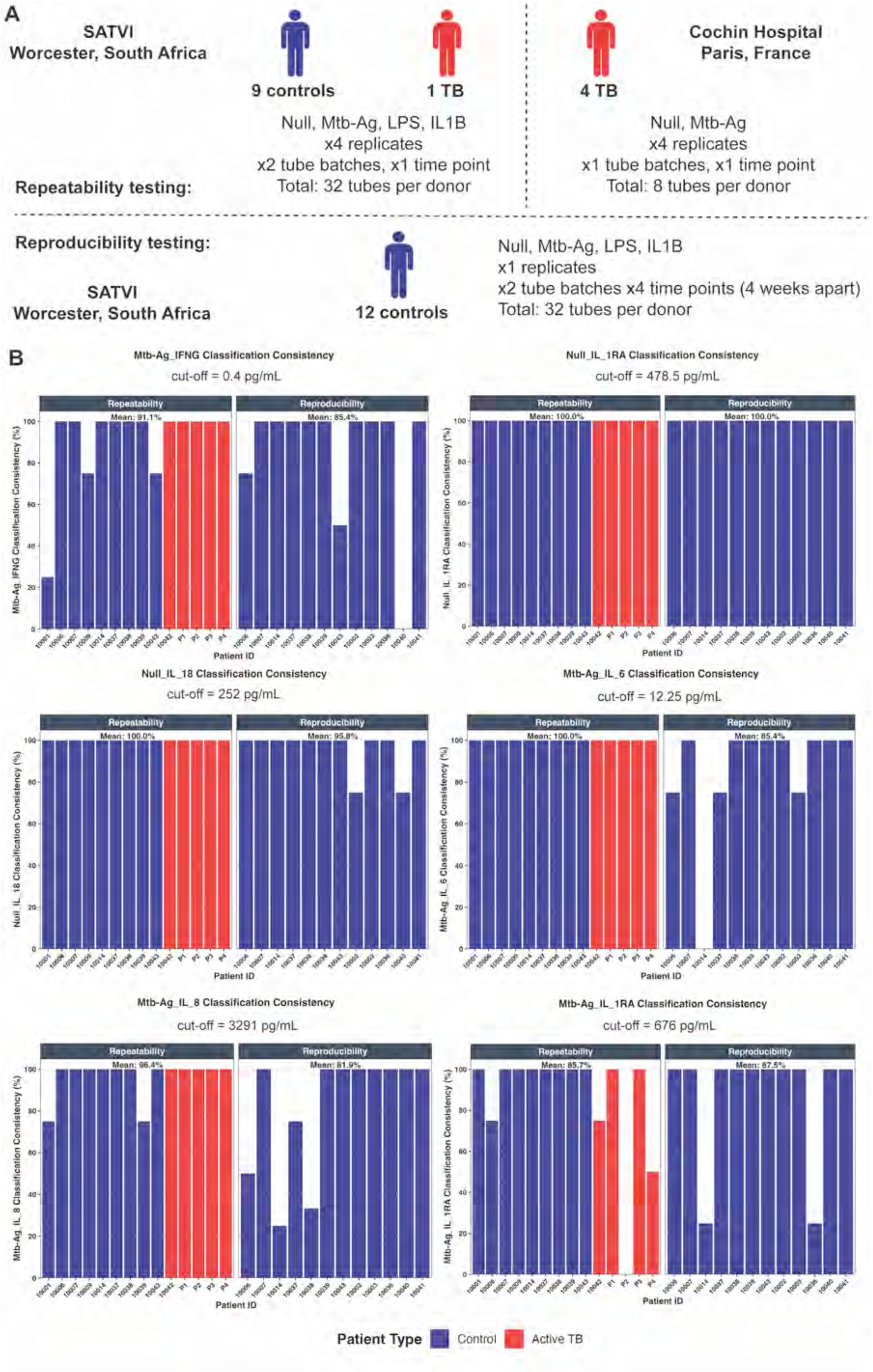
Repeatability and reproducibility testing of TruCulture biomarker based classification for TB diagnosis. (A) Study design for repeatability and reproducibility testing. Controls: asymtomatic QuantiFERON-TB (QFT) positive individuals. (B) Consistency of response in repeatability and reproducibility tests for TruCulture biomarkers to distinguish between controls and active TB patients. Classification formula: Overall consistency = (Absolute (percentage positive - 0.5)/0.5)). Donors 10001 to 10042 were recruited from SATVI in Worcester, South Africa. Donors P1-P4 were recruited from Cochin Hospital in Paris, France.

**Fig. S2.**
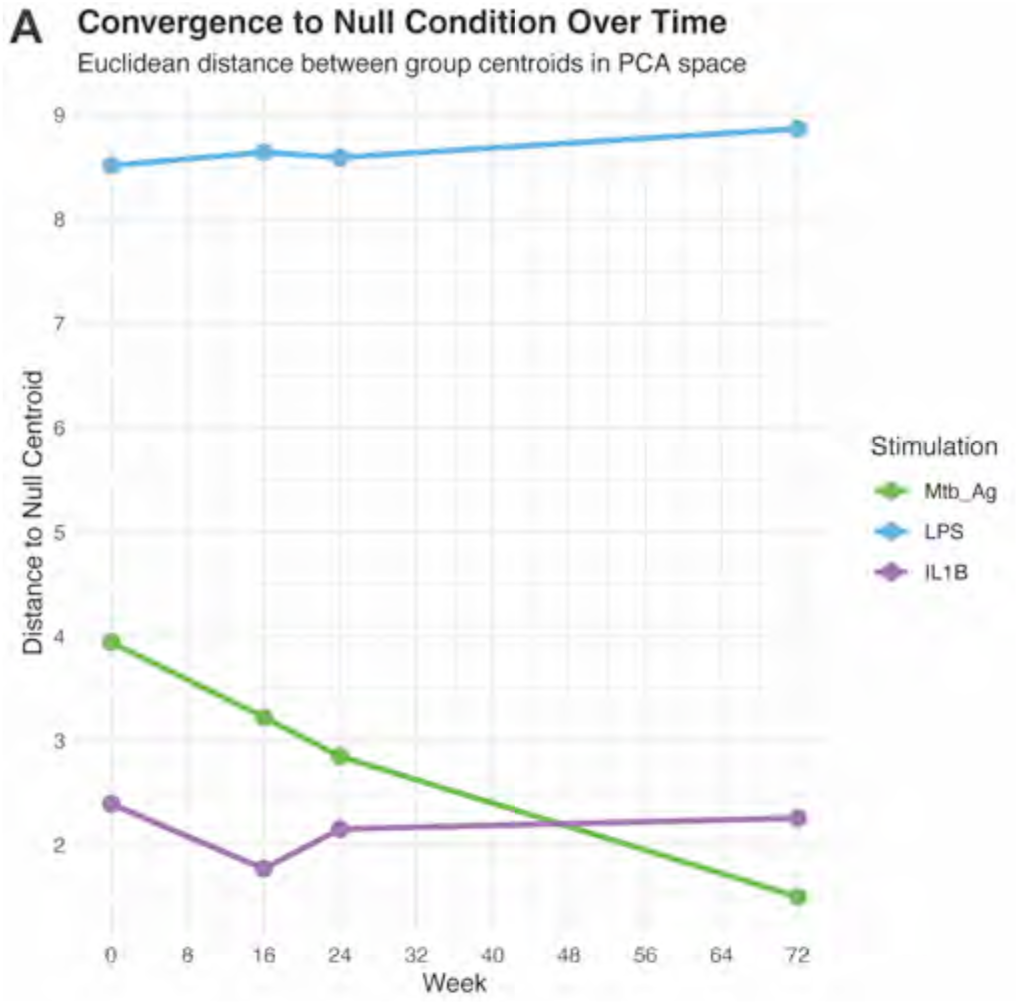
Quantification of PCA cluster distance to Null condition over time. (A) Centroid (Euclidean) distances between Null condition cluster and stimulation clusters before, during, and after TB antibiotic treatment. The distance was computed in the 2D space represented by PC1 and PC2.

**Fig. S3.**
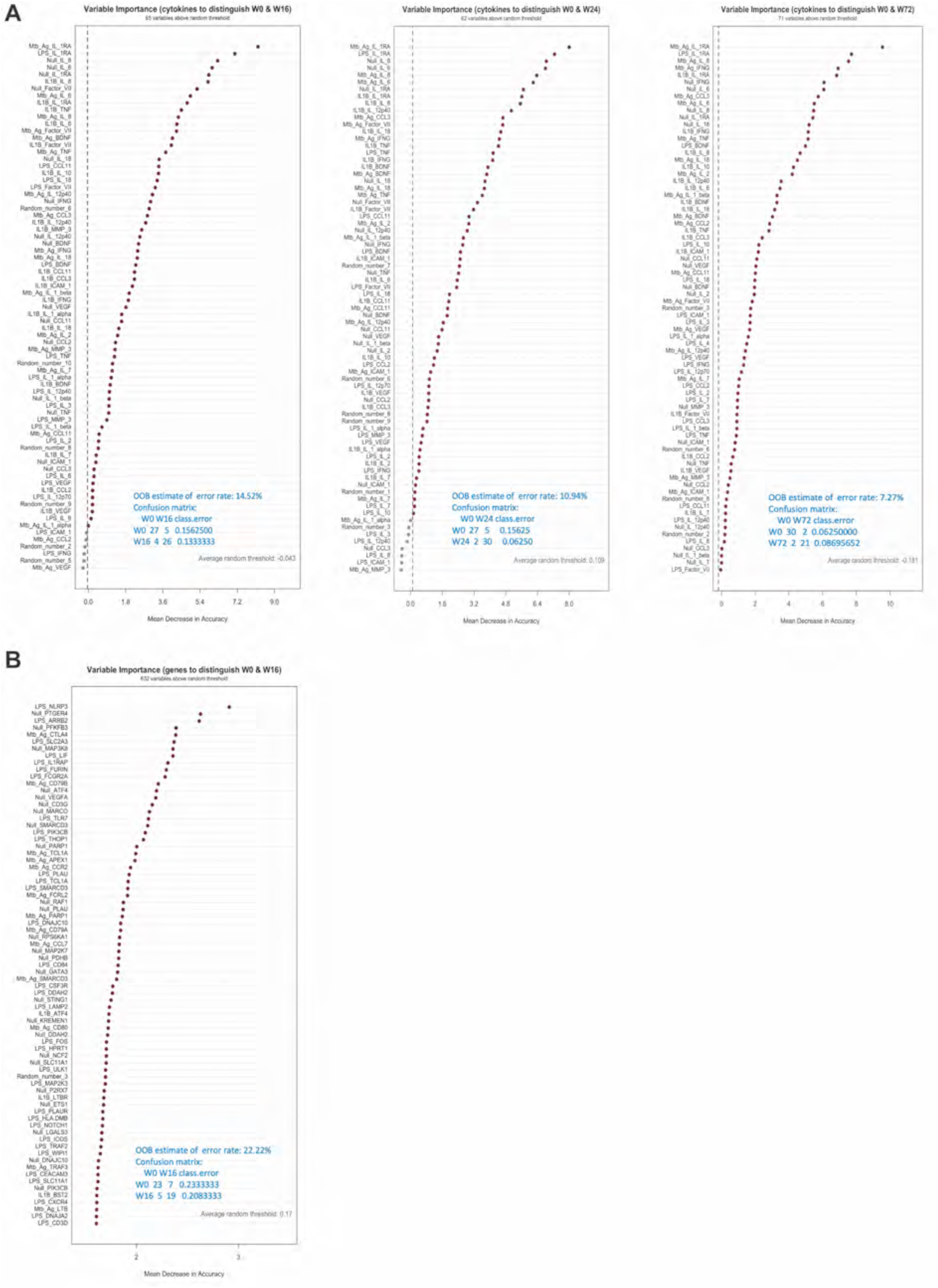
Random forest models to identify biomarkers distinguishing untreated and treated TB patients. (A) Top 75 stimulus_cytokine combinations from random forest models to classify untreated (W0) and treated (W16, W24, and W72) TB patients. (B) Top 75 stimulus_gene expression combinations from random forest model to classify untreated (W0) and treated (W16) TB patients.

**Fig. S4.**
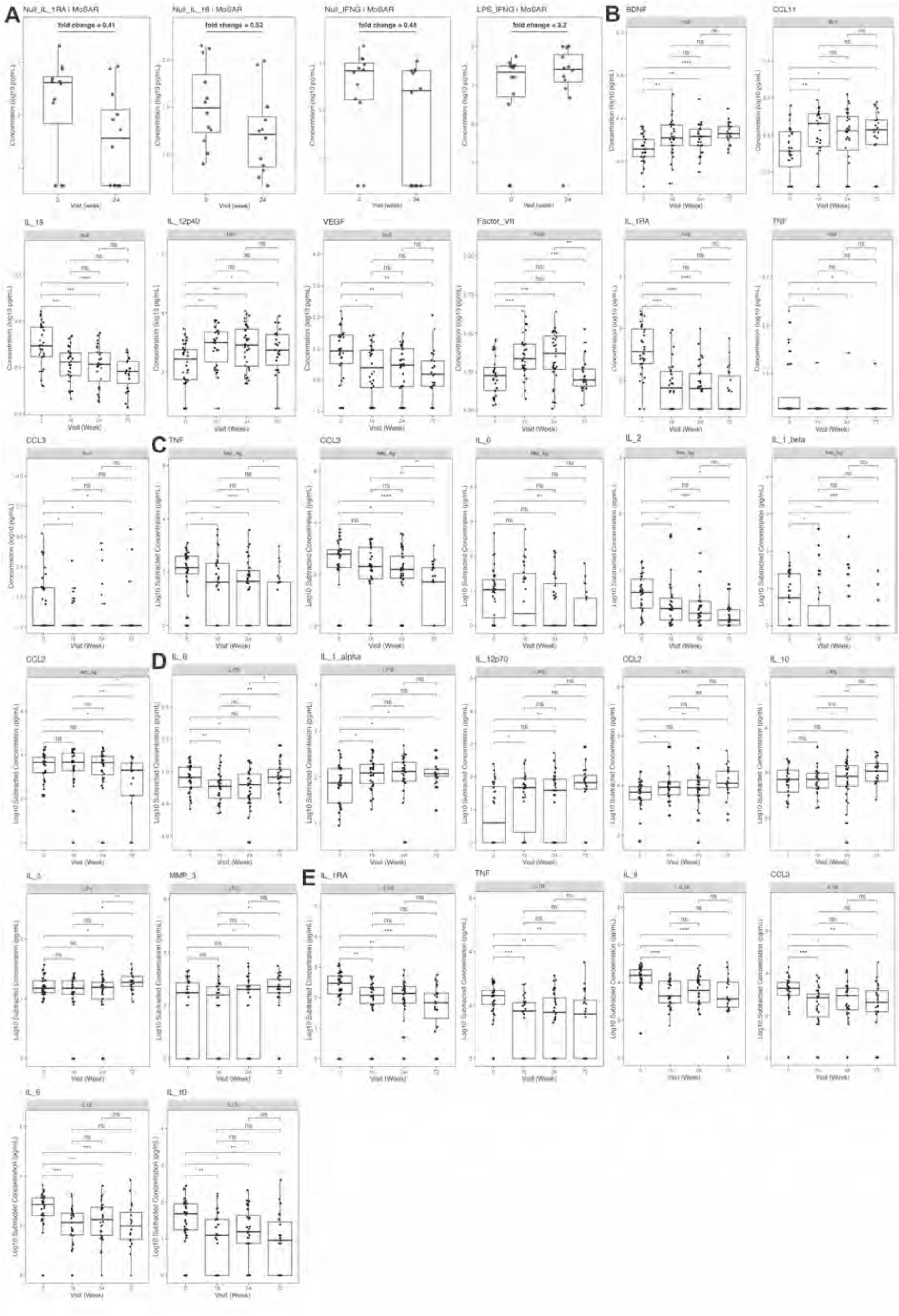
Cytokines for TB disease monitoring. (A) Box plots of cytokine levels showing the median fold change in 12 paired TB patients from an independent replication cohort in Paris (MoSAR), before and after antibiotic treatment. Box plots of cytokine levels showing significant changes during and after successful antibiotic therapy in SATVI cohort under (B) null, (C) *Mtb*-Ag, (D) LPS, and (E) IL-1β stimulation conditions. Cytokine levels in *Mtb*-Ag, LPS and IL1B stimulations are the values subtracted by Null condition. IL-2 concentration was measured by Simoa. P values were calculated using pairwise Wilcoxon tests with FDR adjustment. ns, not significant; *P < 0.05; **P < 0.01; ***P < 0.001; ****P < 0.0001.

**Fig. S5.**
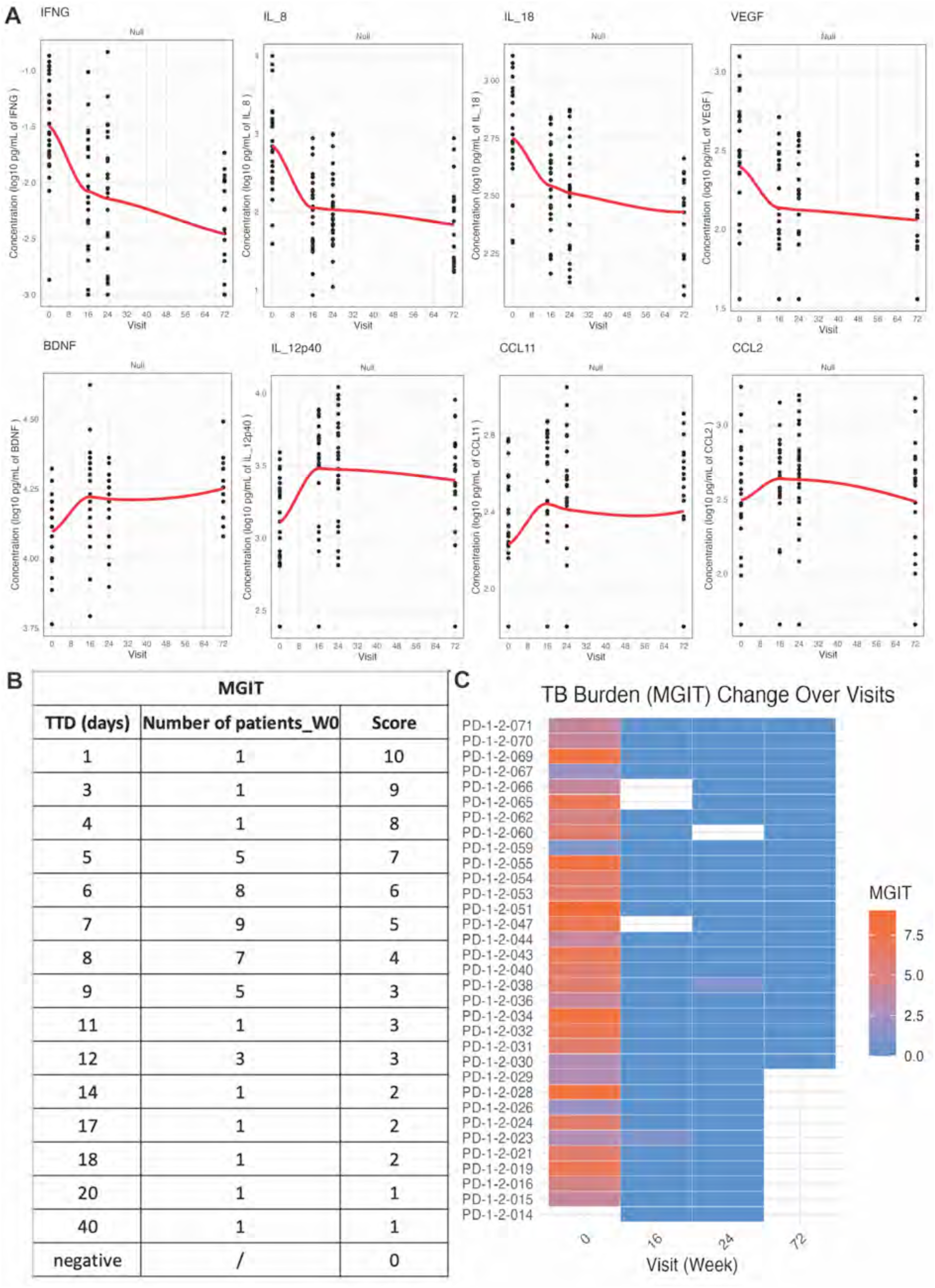
Cytokine responses and sputum *M. tuberculosis* burden kinetics in treatment successful group. (A) Loess regression for cytokine response kinetics under Null condition in patients received 24-week treatment (Arm A and B). (B) Sputum *M. tuberculosis* burden was assessed using time to detection (TTD) from Mycobacteria Growth Indicator Tube (MGIT) liquid culture. MGIT scores were inversely transformed from TTD values to ensure positive correlation with bacterial burden for intuitive interpretation. (C) MGIT measurement showing changes in sputum *M. tuberculosis* burden as TB patients recovered from successful antibiotic therapy.

**Fig. S6.**
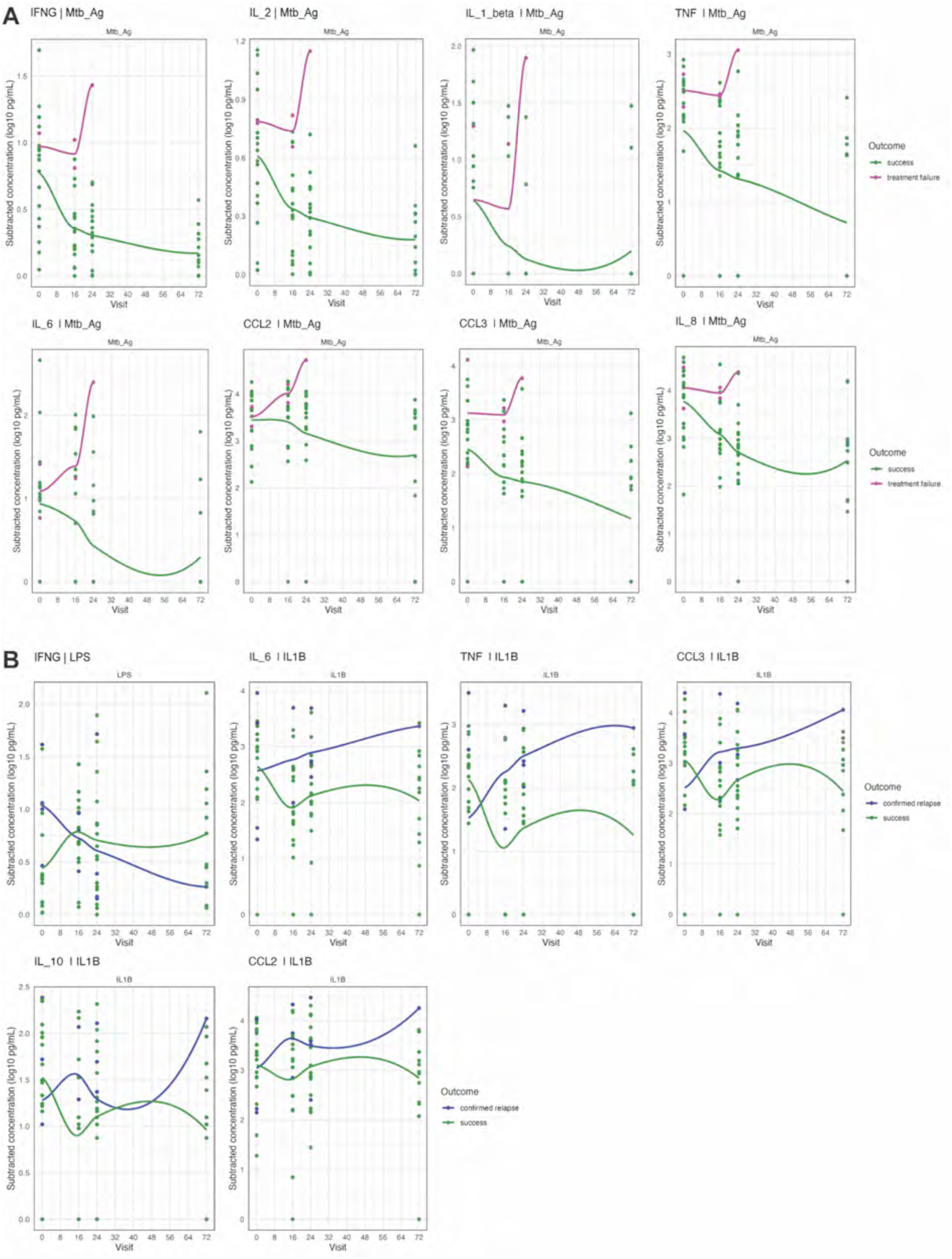
Cytokine response dynamics in different treatment outcomes. (A) Loess regression for cytokine response dynamics in treatment successful group (n=18) and treatment failure group (n=2) in patients with similar lung pathology at baseline (Arm B and C). (B) Loess regression for cytokine response dynamics in treatment successful group (n=18) and confirmed relapse group (n=4) in Arm B and C.

**Fig. S7.**
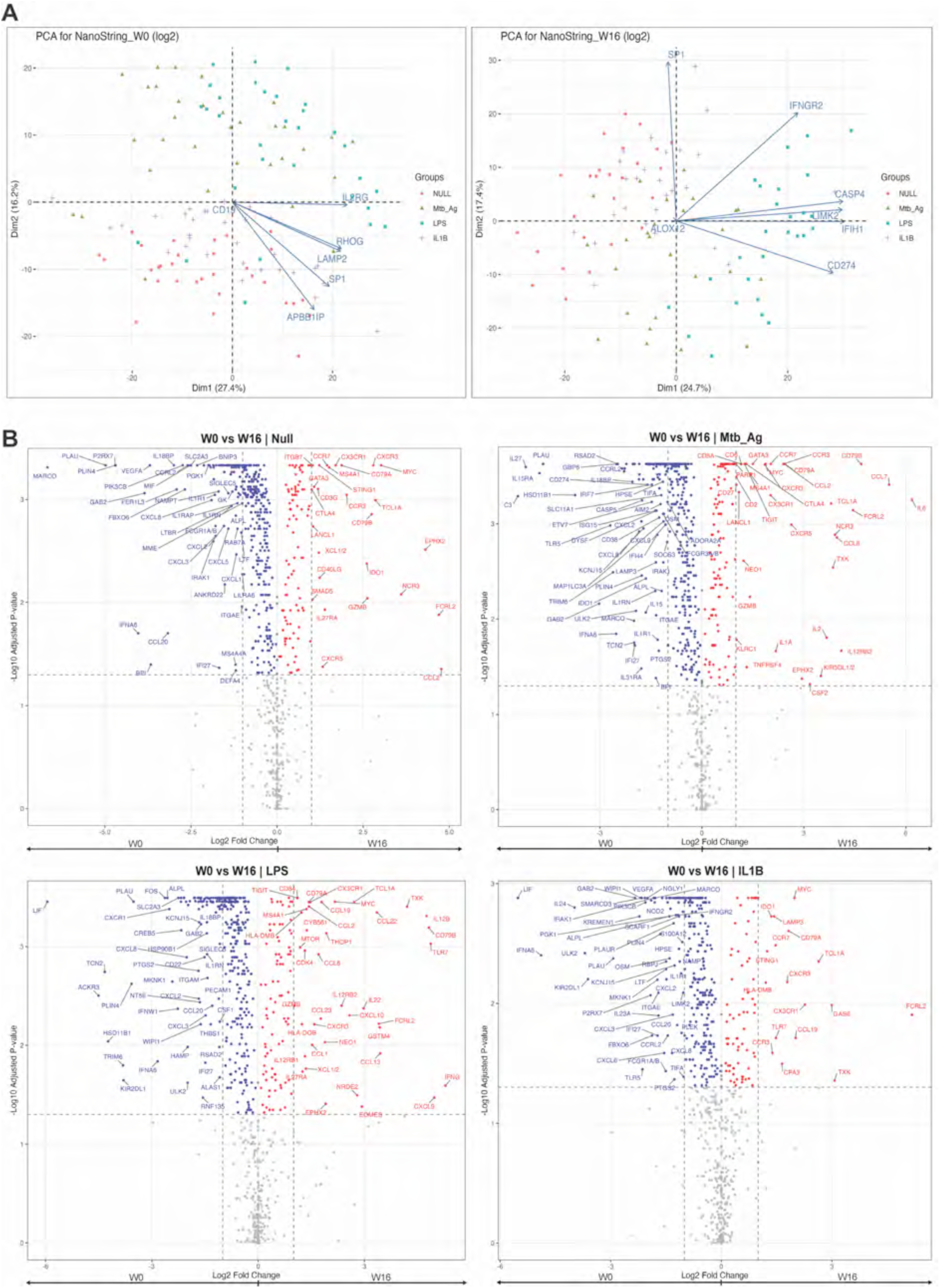
Immune gene expression in untreated and treated TB patients. (A) PCA of the NanoString immune gene expression profiles under four TruCulture conditions (Null, *Mtb*-Ag, LPS, and IL-1β) in untreated (W0) and treated (W16) TB patients. (B) Volcano plots of differentially expressed genes (DEGs) between untreated (W0) and treated (W16) TB patients under null, *Mtb*-Ag, LPS, and IL-1β conditions. P values were calculated using paired Wilcoxon test with FDR adjustment. Positive values (red) are the gene expressions higher in W16; Negative values (blue) are the gene expressions higher in W0.

**Fig. S8.**
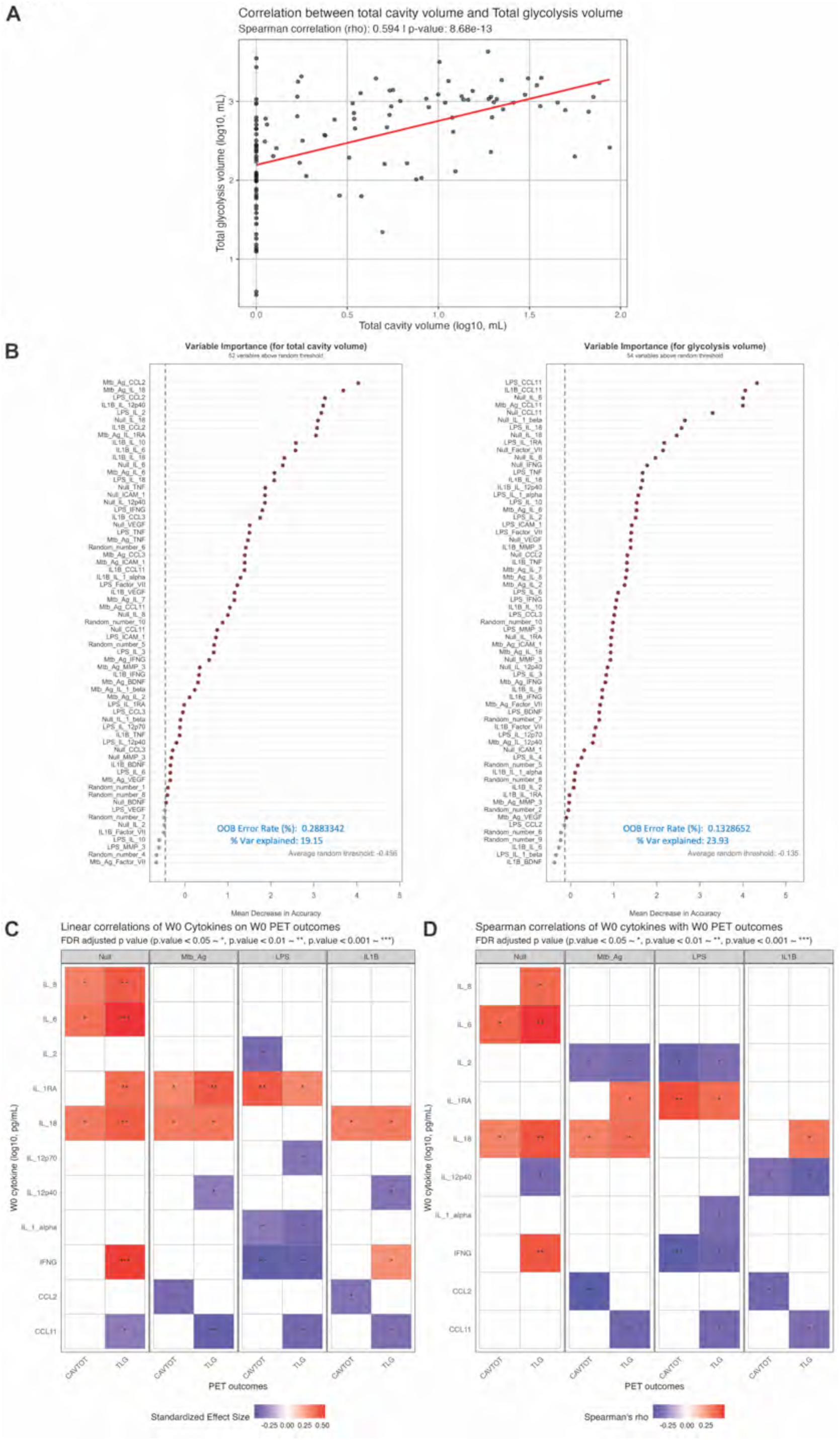
Integration of PET/CT results with cytokine results. (A) Spearman correlation between total cavity volume and total lesion glycolysis volume. (B) Top 65 stimulus_cytokine combinations from random forest models that are associated with total cavity volume and total lesion glycolysis volume in untreated (W0) TB patients. (C) Linear models identifying cytokines significantly associated with total cavity volume (CAVTOT) and total lesion glycolysis volume (TLG) across different stimulation conditions. (D) Spearman correlations identifying cytokines significantly associated with total cavity volume (CAVTOT) and total lesion glycolysis volume (TLG) across different stimulation conditions.

**Fig. S9.**
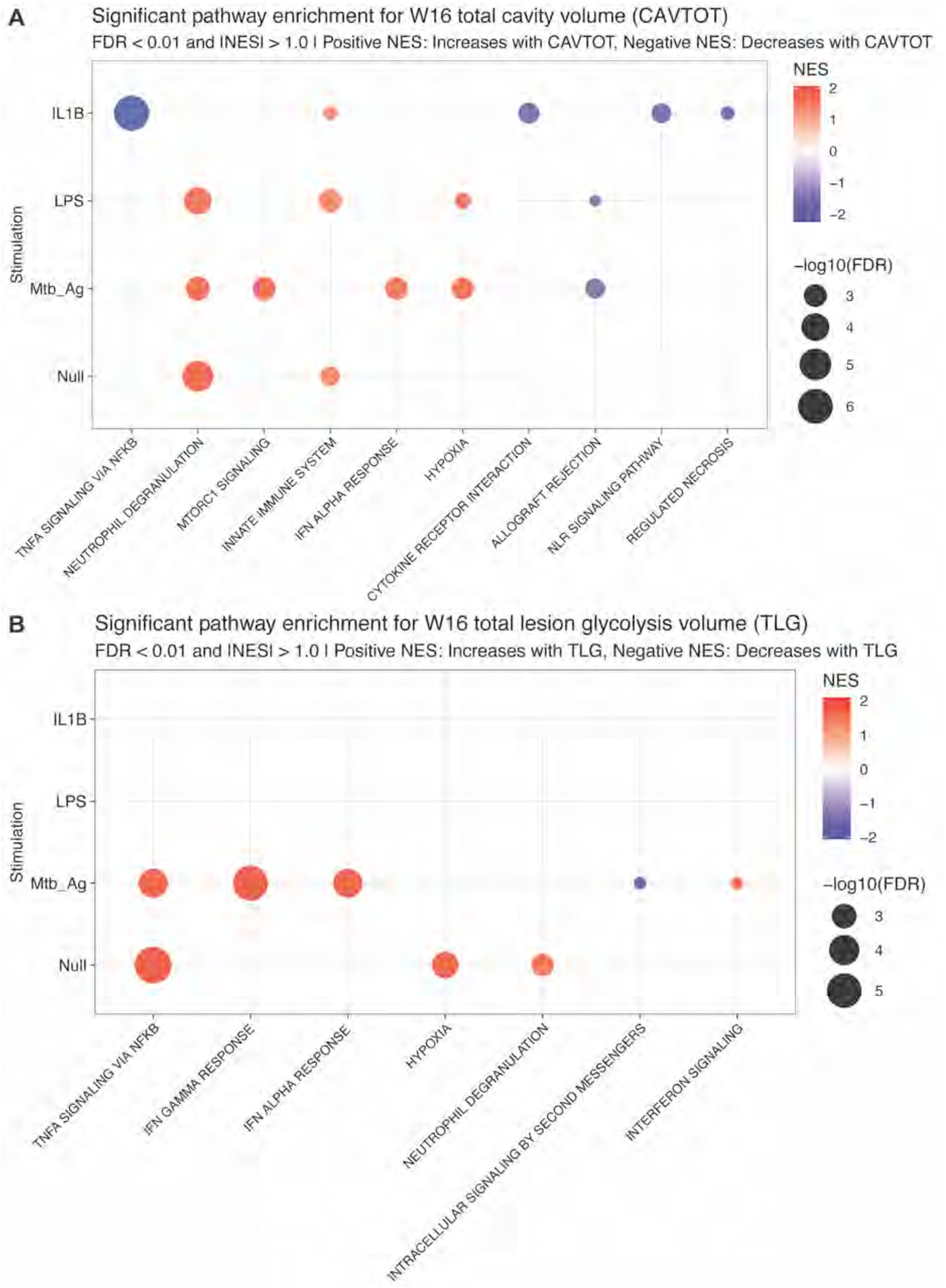
Lung lesion gene expression signatures at week 16. Bubble plot of gene set enrichment analysis showing gene signatures significantly associated with (A) total cavity volume (CAVTOT) and (B) total lesion glycolysis volume (TLG) in treated (W16) TB patients. Bubble color represents the normalized enrichment score (NES), and bubble size represents FDR-adjusted P values.

**Fig. S10.**
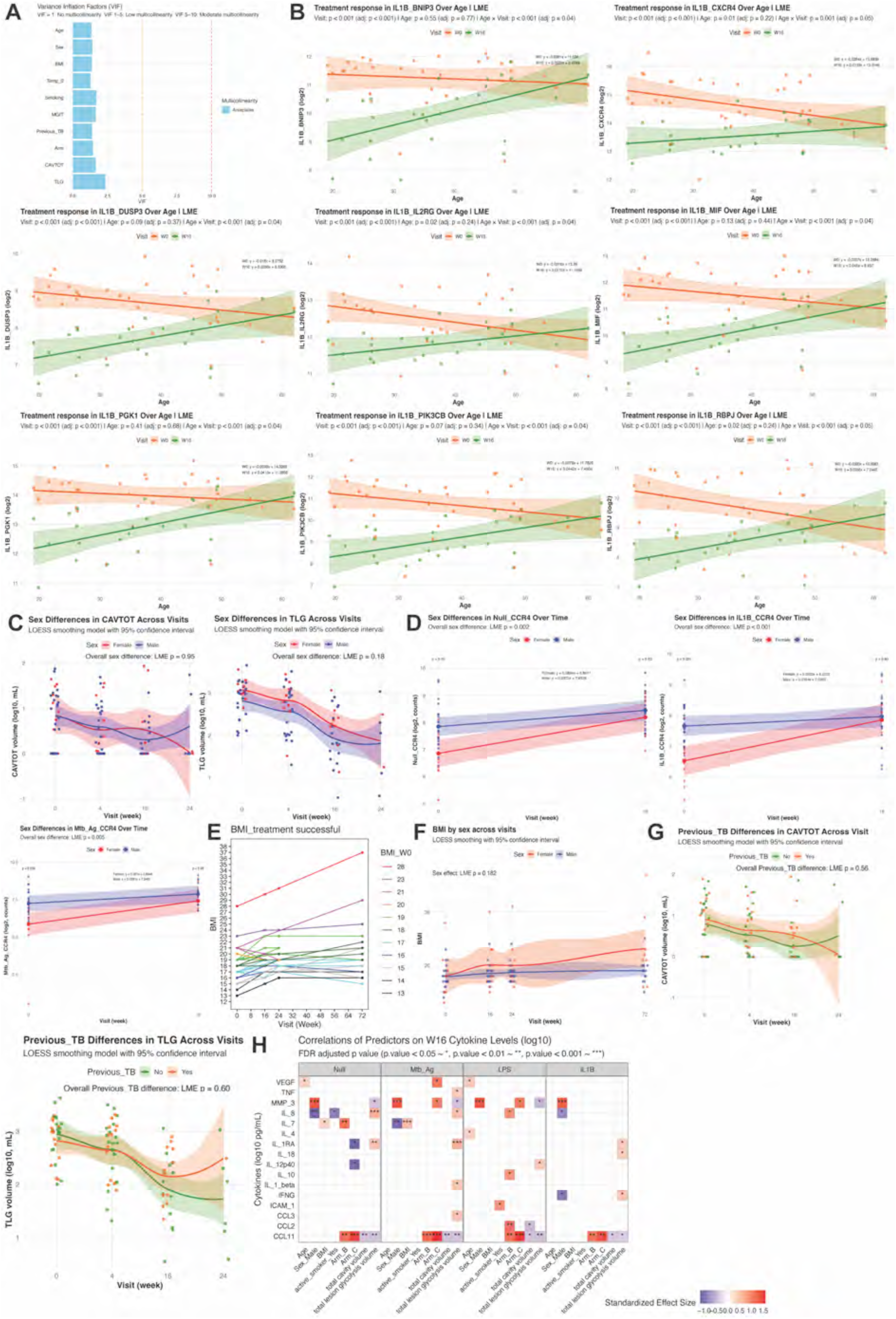
**Impact of demographic and clinical factors on TB patients**. (A) Assessment of multicollinearity among demographic and clinical factors using the variance inflation factor (VIF). (B) Linear mixed-effects (LME) models of significant age and treatment effects on NanoString gene expression responses, with 95% confidence interval. (C) LOESS smoothing of sex and treatment effects on total cavity volume (CAVTOT) and total lesion glycolysis volume (TLG) throughout successful TB treatment. (D) Linear mixed-effects models of significant sex and treatment effects on NanoString gene expression responses, with 95% confidence interval. (E) BMI change of TB patients throughout and after successful antibiotic therapy. Line colours represent BMI value before treatment at W0. (F) BMI distributions in female and male before, during and after TB treatment. (G) LOESS smoothing of previous TB and treatment effects on total cavity volume (CAVTOT) and total lesion glycolysis volume (TLG) throughout successful TB treatment. (H) Linear models identifying demographic and clinical factors significantly associated with W16 cytokine levels across different stimulation conditions. Body temperature and *M. tuberculosis* burden were excluded from week 16 linear model due to minimal variance following TB treatment.

**Fig. S11.**
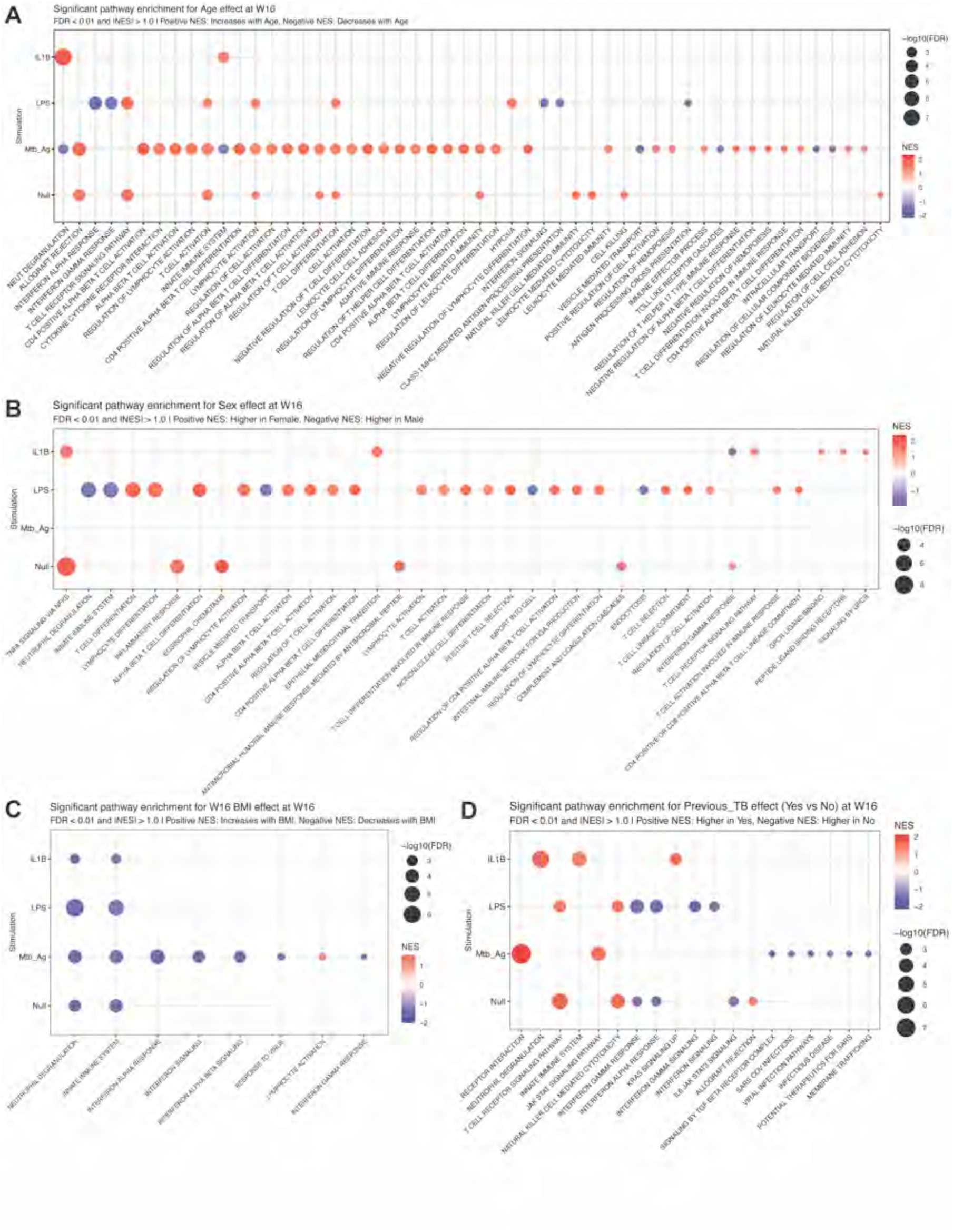
Week 16 gene signatures associated with demographic and clinical factors. Bubble plot of gene set enrichment analysis showing gene signatures significantly associated with (A) age, (B) sex, (C) BMI and (D) previous TB in treated (W16) TB patients. Bubble color represents the normalized enrichment score (NES), and bubble size represents FDR-adjusted P values.

**Table S1. GSEA significant pathways and leading-edge genes**

(Supplemental excel file)

## Notes

### Competing Interest Statement

The authors have declared no competing interest.

